# Engineering microscale systems for fully autonomous intracellular neural interfaces

**DOI:** 10.1101/689562

**Authors:** Swathy Sampath Kumar, Michael S. Baker, Murat Okandan, Jit Muthuswamy

## Abstract

Conventional electrodes and associated positioning systems for intracellular recording from single neurons *in vitro* and *in vivo* are large and bulky, which has largely limited their scalability. Further, acquiring successful intracellular recordings is very tedious, requiring a high degree of skill not readily achieved in a typical laboratory. We report here a robotic, MEMS-based intracellular recording system to overcome the above limitations associated with form-factor, scalability and highly skilled and tedious manual operations required for intracellular recordings. This system combines three distinct technologies: 1) novel microscale, glass-polysilicon penetrating electrode for intracellular recording, 2) electrothermal microactuators for precise microscale movement of each electrode and 3) closed-loop control algorithm for autonomous positioning of electrode inside single neurons. Here, we demonstrate the novel, fully integrated system of glass-polysilicon microelectrode, microscale actuators and controller for autonomous intracellular recordings from single neurons in the abdominal ganglion of *Aplysia Californica* (n = 5 cells). Consistent resting potentials (< −35 mV) and action potentials (> 60 mV) were recorded after each successful penetration attempt with the controller and microactuated glass-polysilicon microelectrodes. The success rate of penetration and quality of intracellular recordings achieved using electrothermal microactuators were comparable to that of conventional positioning systems. The MEMS-based system offers significant advantages: 1) reduction in overall size for potential use in behaving animals, 2) scalable approach to potentially realize multi-channel recordings and 3) a viable method to fully automate measurement of intracellular recordings. This system will be evaluated in vivo in future rodent studies.

## Introduction

Intracellular recordings from single neurons provide functional information at the highest spatial and temporal resolution among known techniques for brain monitoring. They offer several significant advantages over extracellular recordings: 1) ability to record sub-threshold dynamic events such as synaptic potentials and membrane potential oscillations, which have been identified to play important roles in neural coding [1]–[4], 2) large dynamic range of signal (80-100 mV) compared to signal recorded with extracellular electrodes (hundreds of μV to 1 mV) and 3) ability to obtain structural information (through dye labeling, passive membrane properties) about the neurons being recorded from in vivo, thereby allowing correlation of structure with function at single neuron resolution. A more recent alternative approach to intracellular recording is voltage imaging with genetically encoded voltage indicators (GEVIs). Although current GEVIs can image sub and supra threshold events from neuronal ensembles at single neuron resolution, they do not match the temporal resolution and sensitivity of the intracellular recording technique [5]. Thus, the quality of information obtained with intracellular recordings is unparalleled and fundamental to our understanding of neural computation and function.

Traditionally, glass micropipettes integrated with cumbersome micromanipulators and bulky positioning systems have been used to record intracellularly. Although they have been extensively used for in vitro studies, their use in in vivo studies has been limited due to their large form factor. The conventional intracellular recording system has significant limitations: 1) due to the large form factor of the technologies involved, recordings have mostly been obtained from anesthetized animals with the exception of a few studies [6], [7], 2) recordings are obtained from one neuron at a time (serial recording), 3) their use requires extraordinary manual skill and tedious operations, which results in a long training period for neurophysiologists, 4) duration of recordings obtained are typically short (45-60 min in anesthetized animals, 5-30 min in awake head-fixed animals [8], [9]), often due to mechanical disruptions at the electrode-cell interface. These challenges have impeded chronic intracellular recording studies from a population of neurons in anesthetized and awake animals. The ability to record intracellularly from neuronal networks in freely behaving animals would allow correlation of ultra-high resolution functional information with modulative behavior and accelerate neurophysiological studies on mechanisms of neuronal function and dysfunction.

Several approaches have been reported recently to address some of the limitations of manual operations in the conventional intracellular recording system. Recently, automated systems have been developed to reduce the ‘art’ in the process of intracellular recording in vivo [10]–[12]. Kodandaramiah and colleagues [10] developed a closed-loop control system that used a temporal sequence of electrode impedance changes as a feedback signal to automate movement of electrode and whole-cell patching of neurons in cortex and hippocampus of anesthetized, head-fixed mice. They recently improved the algorithm to automate localization of pipette to deep cortical nuclei through autonomous detection and lateral navigation around blood vessels and obtained high-yield (10%) thalamic whole cell recordings [13]. Desai et al.[12] and Ota et al.[11] developed similar algorithms to automate cortical whole-cell patching in awake, head-fixed, behaving mice and sharp micropipette recording in anesthetized, head-fixed mice respectively.

Conventional techniques to achieve long-duration intracellular recordings in vivo include draining of cerebrospinal fluid, rigid fixation of cranium to recording apparatus and passive stabilization using floating micropipettes [14]. Fee [15] developed a novel control strategy that compensated for residual brain motion in awake, head-fixed rats due to cardiac and respiratory pulsations as well as spontaneous movement of animal. Through dynamic stabilization of a sharp micropipette relative to the brain, intracellular recordings were obtained for ∼9 min from resting rats (< 10 s with no active stabilization) and ∼3 min from active rats. Although successful, all the above technologies used bulky microdrive systems and glass micropipettes, which prevented their immediate translation to freely behaving animals and feasibility of parallel intracellular recordings.

In the first of its kind, Lee et al [16] introduced a technique to record intracellularly from motor cortex and hippocampus of non-head-fixed, freely moving rats. They developed and used a head-mounted device consisting of a miniaturized recording headstage and a miniaturized motor integrated with a patch pipette holder. Mechanical stabilization was achieved by anchoring the recording pipette to the skull using dental acrylic after establishing a whole-cell recording [16], [17]). Recently, they extended the technique to mice by using a UV- transparent collar and a UV cured adhesive for pipette fixation[18]. Long et al [4] developed a miniaturized linear microdrive to record intracellularly from freely moving song birds using sharp micropipettes. Although these groups recorded for long durations (mean recording time of 10 min) in freely behaving animals, use of glass micropipettes limited recording to one neuron per animal. Further, there was no mechanism to reposition electrode upon loss of recordings.

Several groups have developed novel, metal-based, micro/nanoscale electrodes to potentially realize multi-channel intracellular recordings [19]–[23]. These electrodes successfully recorded synaptic and intracellular-like or full-blown action potentials in neuronal cultures and brain slices, however they have not been demonstrated in vivo. Recently, Moore et al.[24] showed intracellular-like signals recorded from dendritic arbors in cortex of freely moving rats using chronically implanted tetrodes. Although they recorded intracellular-like signals for several hours to days for the first time, the success-rate of the technique (13%) was relatively low.

Therefore, there is a critical need for a technology that enables multi-channel intracellular recordings from unrestrained, behaving animals, which is currently unavailable. We report here a novel microscale, robotic, intracellular positioning and recording system as a first step towards addressing the above need. We report successful demonstration of (1) MEMS-based technologies to significantly reduce the form factor of the recording electrode as well as the positioning system required to move the intracellular electrodes, thereby addressing the size and scalability challenges of the conventional electrode navigation systems and (2) closed loop control technology to automate electrode movement to seek and penetrate neurons, and maintain intracellular recordings that will minimize the training barriers for personnel using such systems. In this study, we demonstrate the ability of this miniaturized MEMS based system to autonomously isolate, impale and record intracellular signals from single neurons in the abdominal ganglion of *Aplysia Californica*. Future studies will test this system in vivo.

### Microscale intracellular recording system

The MEMS sub-systems of the proposed intracellular recording system are illustrated in Fig. 1. The two key sub-systems are: (a) glass-polysilicon microelectrode - polysilicon microelectrode integrated with a miniaturized glass micropipette to penetrate and record intracellular potentials from single neurons (Fig. 1A & 1C) and (b) electrothermal microactuators for precise microscale navigation and positioning of glass-polysilicon microelectrode inside single neurons (Fig. 1A & 1B). This system is integrated with a closed-loop control algorithm to enable autonomous movement of microelectrode, isolation and penetration of neurons (Fig. 8).

**Fig.1:**
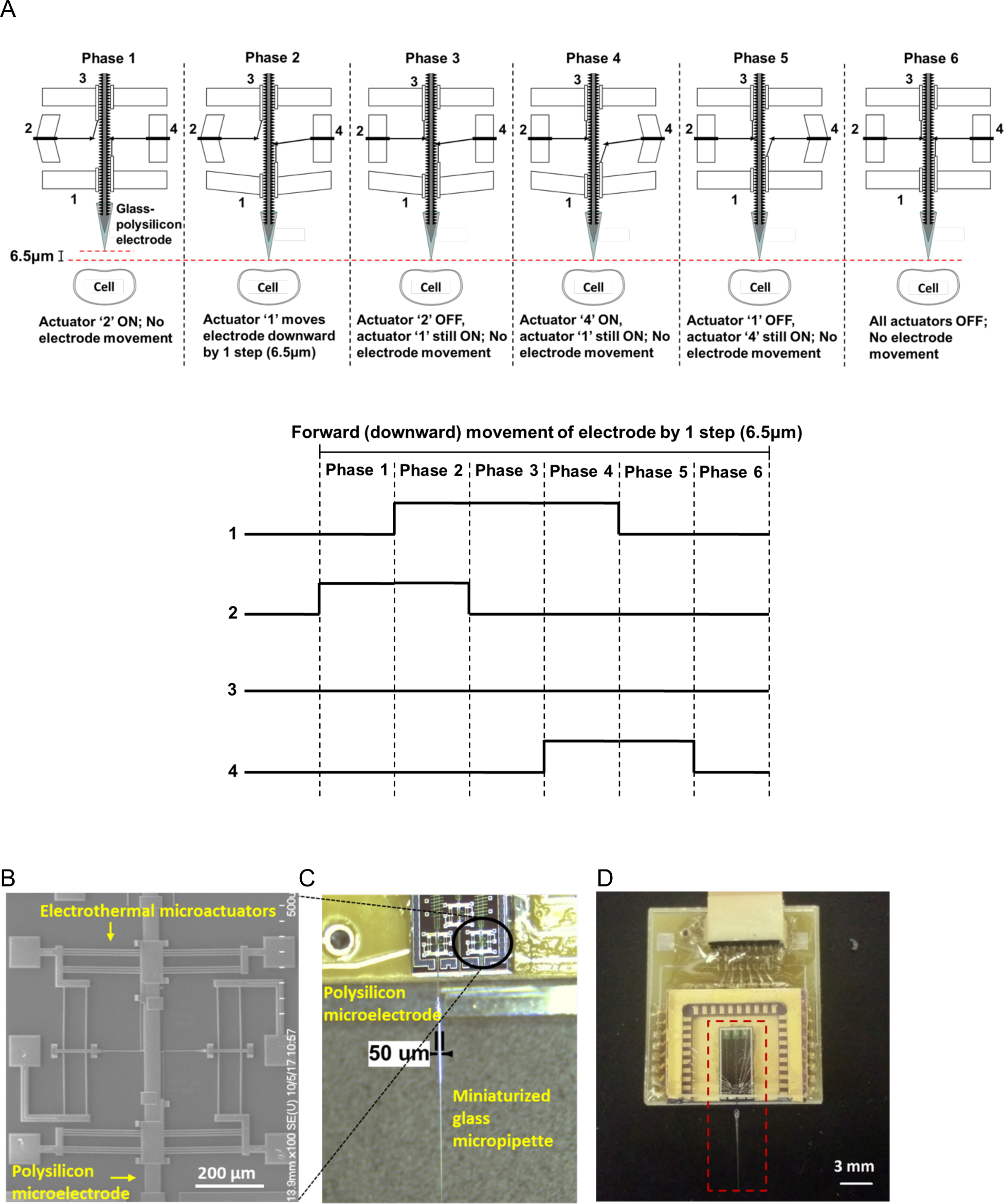
MEMS-based system for microscale actuation and intracellular recording. (A) Mechanism of actuation using chevron-peg electrothermal microactuators - 6 distinct phases in the forward (downward) actuation towards a neuron by 1 step (6.5 μm) using the 4 different electrothermal microactuators (1 - Forward drive, 2 - Disengage reverse, 3 - Reverse drive, 4 - Disengage forward). Their corresponding voltage waveforms are shown below. (B) Micrograph of the electrothermal microactuators integrated with the polysilicon microelectrode to enable cell penetration. (C) Integrated glass-polysilicon microelectrode for intracellular recording. (D) The integrated MEMS-based intracellular recording system.

### Principle of actuation of the electrothermal microactuators

Bi-directional movement of each glass-polysilicon microelectrode is enabled by the chevron-peg mechanism reported earlier [25]. The mechanism of forward (or downward) actuation of the electrode is shown in Fig. 1A. Briefly, the polysilicon microelectrode has a set of teeth spaced 6.5 μm apart on both sides along the length of the electrode. A peg/pawl engages the teeth and holds the microelectrode in position during rest conditions. Each microelectrode is coupled with two pairs of electrothermal actuators. The ‘forward drive’, working in conjunction with ‘disengage forward’ and ‘disengage reverse’ actuators enables forward (downward) movement, while the ‘reverse drive’, working in conjunction with ‘disengage forward’ and ‘disengage reverse’ actuators enables reverse (upward) movement of the microelectrode. Each actuator is composed of an array of doped polysilicon beams anchored at two ends and attached to a central shuttle as shown on either side of the polysilicon microelectrode in Fig. 1B. Application of voltage pulses typically 6-10 V amplitude causes thermal expansion of the beams, which causes displacement of the shuttle. The central shuttles of the drive and disengage actuators are both connected to a peg in an L-shaped arrangement to facilitate movement. A pre-programmed set of pulsed voltage waveforms applied to these actuators allow movement of the microelectrode in forward/reverse direction. The voltage waveforms for the actuators to enable forward movement of the microelectrode by one step (6.5 μm) are also shown in Fig. 1A. Details on optimal parameters for reliable activation of the microactuators and the microstructural details of the assembly can be found in our prior report [25].

## Results

### Intracellular recordings using glass-polysilicon (GP) microelectrode

In this study, we report a novel glass-polysilicon (GP) microelectrode for intracellular recording. The GP microelectrodes consistently recorded good quality resting potentials (RP < −35 mV) and/or action potentials (AP >70 mV) from neurons in the abdominal ganglion of *Aplysia Californica*, similar to conventional glass micropipettes (Fig. 2A). The quality of signals recorded with the GP microelectrode from abdominal ganglion neurons was comparable to recordings acquired using conventional glass micropipettes, as shown in Fig. 2B (from n = 3 distinct GP microelectrodes). Unpaired *t*-test comparing the means of signal (RP and peak-peak AP) amplitudes recorded with the two electrodes showed no statistically significant difference between them (RP: *p*>0.5 and AP: *p*>0.2). The GP microelectrodes also recorded good quality resting potential (V_m_ = −74 mV) from a cell in the motor cortex of an anesthetized rat at a depth of 800 μm from the surface of the brain (Fig. 2C).

**Fig.2:**
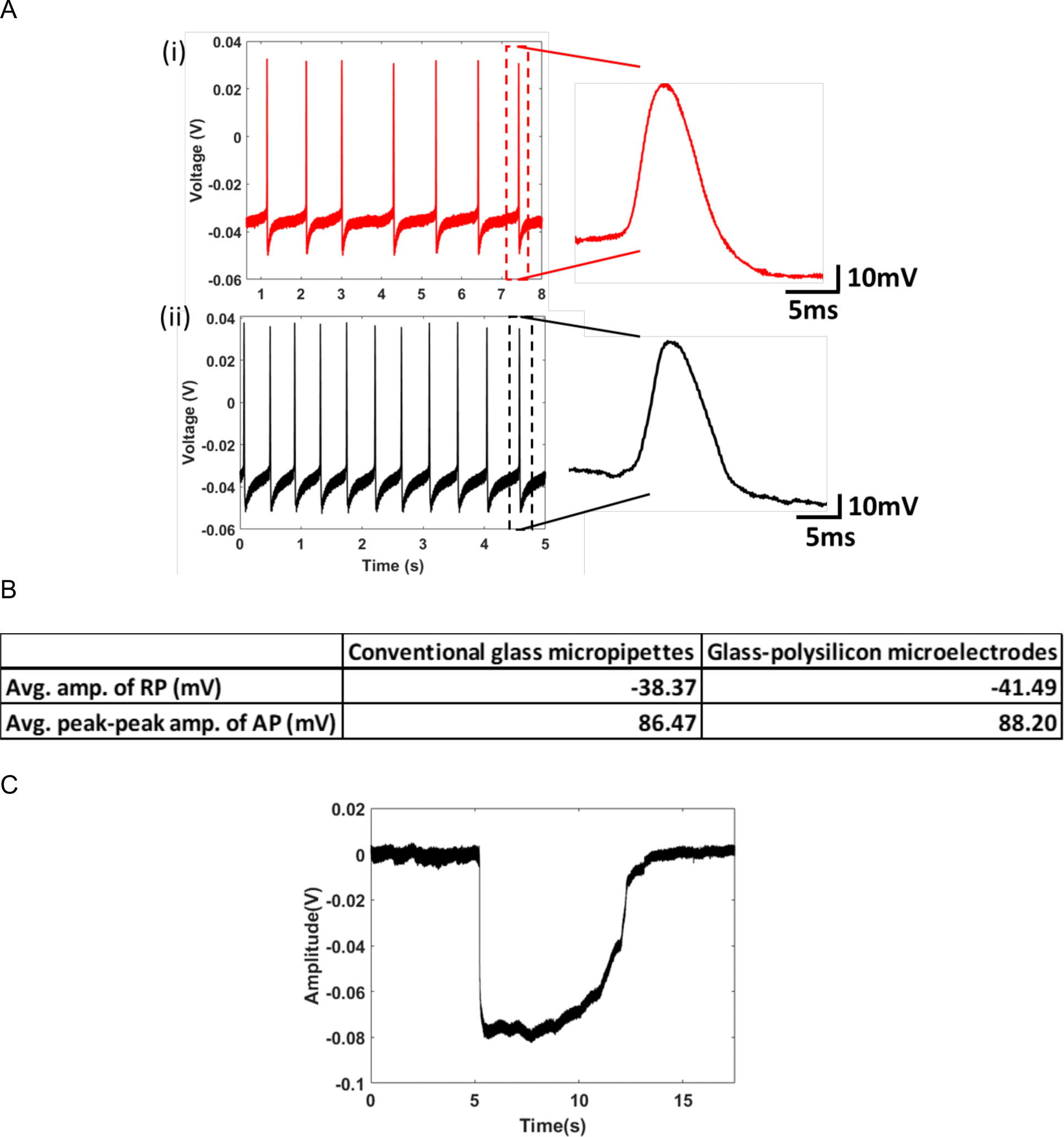
Intracellular recordings using the glass-polysilicon (GP) microelectrode. (A) (i) Intracellular recordings (RP = −35.5 mV, peak-peak AP = 80.1 mV) obtained from an isolated abdominal ganglion neuron in Aplysia using a conventional glass micropipette and (ii) using GP microelectrode (RP = −38.8 mV, peak-peak AP = 85.5 mV). (B) Comparison of the quality of intracellular signals using GP microelectrodes with those obtained using conventional glass micropipettes (n=3 neurons) showing no significant difference (RP: *p*>0.5, AP: *p*>0.2). (C) In vivo intracellular recording (RP = −74 mV) obtained from the motor cortex of a rat using a GP microelectrode. (RP: Resting Potential, AP: Action Potential).

### Electrical impedance of glass-polysilicon (GP) microelectrode

The closed loop control for autonomous isolation and penetration of neurons uses DC electrical impedance of the tip of the electrode and measured voltage at the tip of the electrode as feedback variables (Fig. 8). A model of the glass-polysilicon (GP) microelectrode-neuron interface (Fig. 3A, B) was constructed to predict the electrical impedance of this electrode at DC. All simulations were done using Simulink™ (Mathworks Inc., Natick MA). The following parameters were used for the model: 1) neuronal membrane resistance (*R*_*m*_) of 25 MΩ and membrane capacitance (*C*_*m*_) of 500 pF, obtained from prior studies by Hai et al. [27] and Ungless et al. [28]; 2) Miniaturized pipette tip resistance (*R*_*tip*_) of 30 MΩ and distributed capacitance (*C*_*d*_) of 15 pF, as measured from voltage responses of conventional micropipettes with silver/silver chloride electrode to current pulses of 1 nA (*R*_*tip*_ = steady-state value of voltage response to 1 nA current injection and *C*_*d*_ = measured time-constant/R_tip_); 3) Polysilicon charge transfer resistance (*R*_*ct*_) of 2.35 GΩ, double layer capacitance (*C*_*dl*_) of 3.95 nF and solution resistance (*R*_*s*_) of 16 KΩ, were obtained from the electrochemical impedance spectrum of a polysilicon microelectrode measured with a electrochemical workstation (CH Instruments Inc.) and subsequently modeled using ZSimpWin™ software.

**Fig.3:**
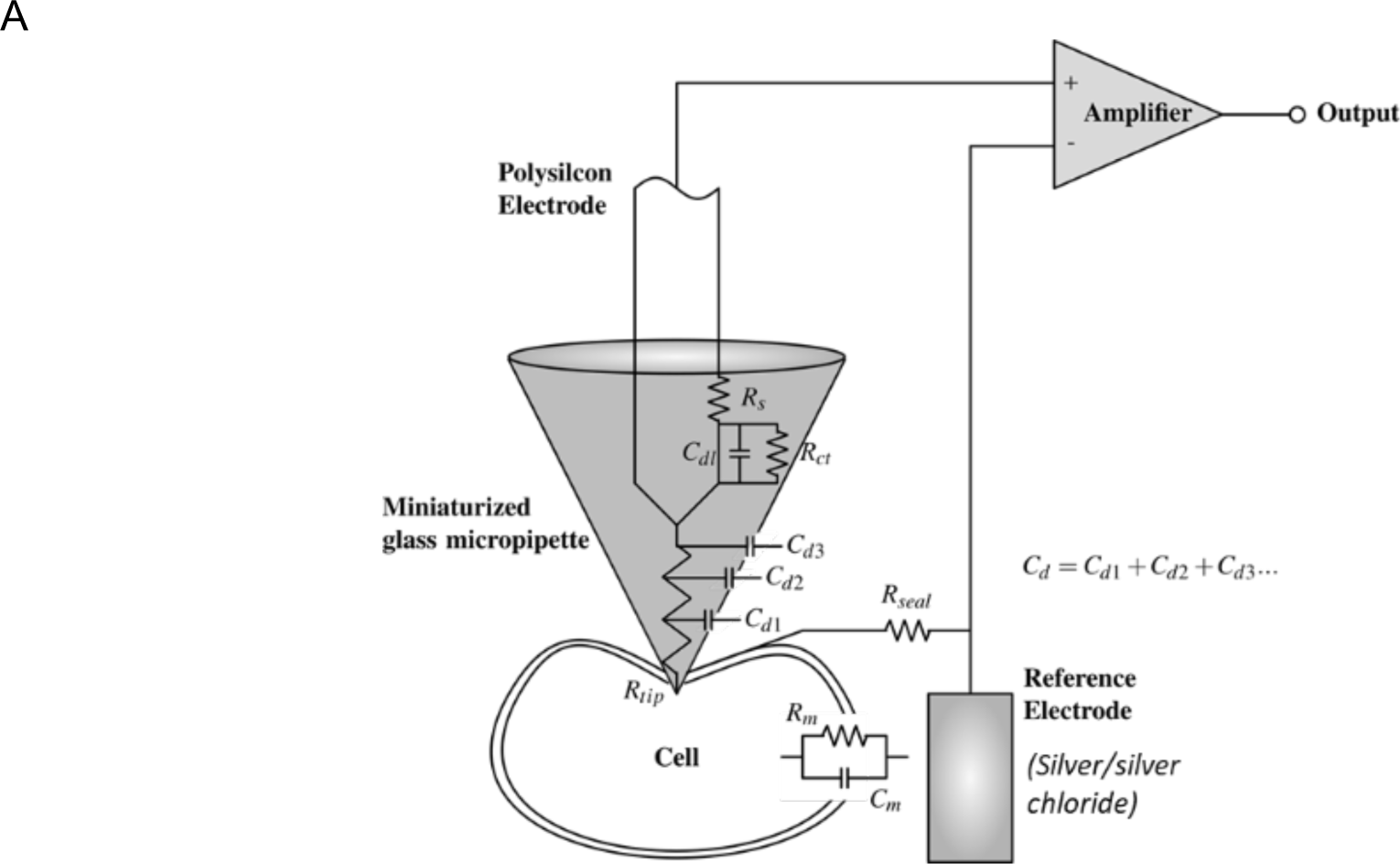
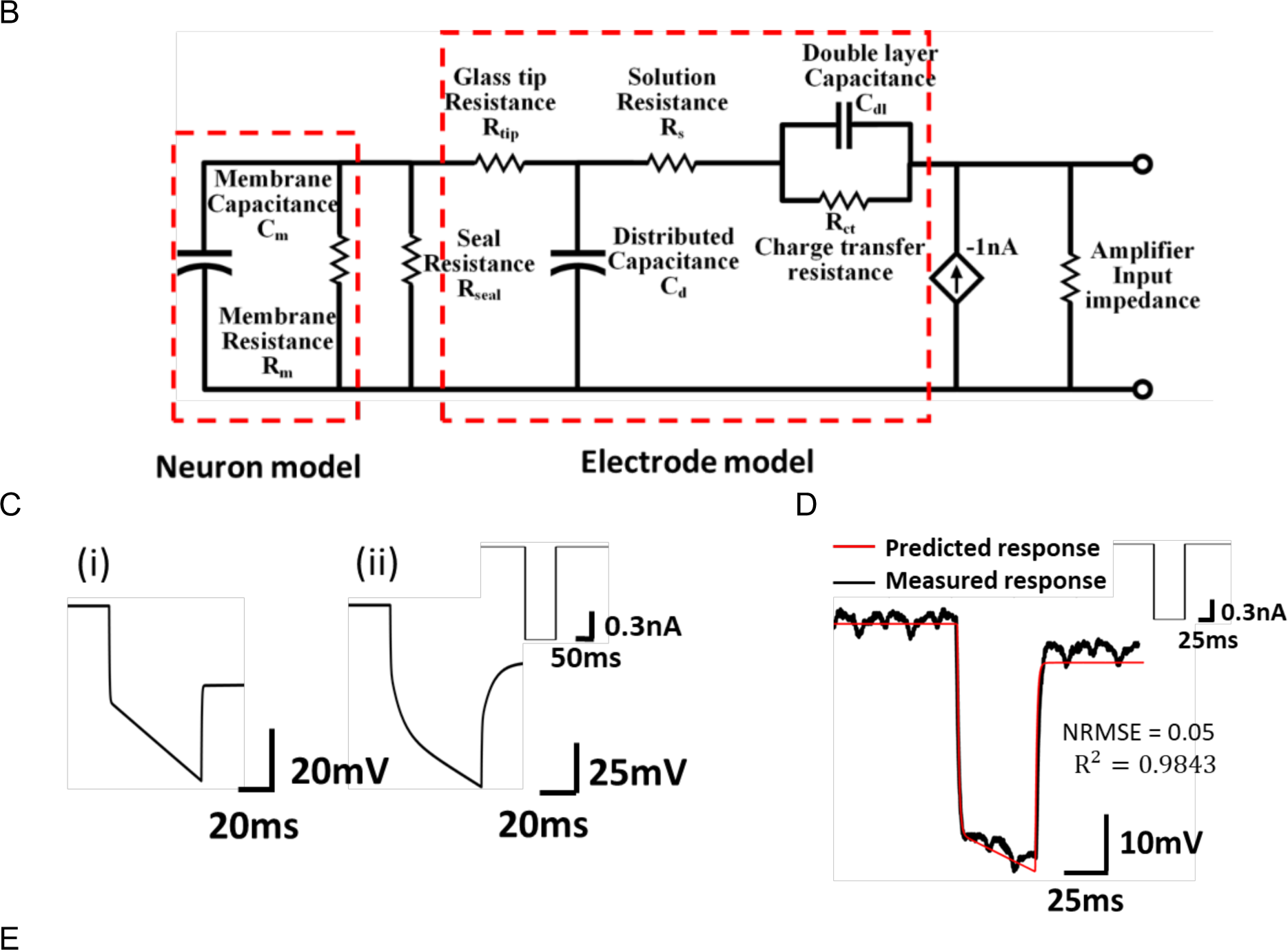
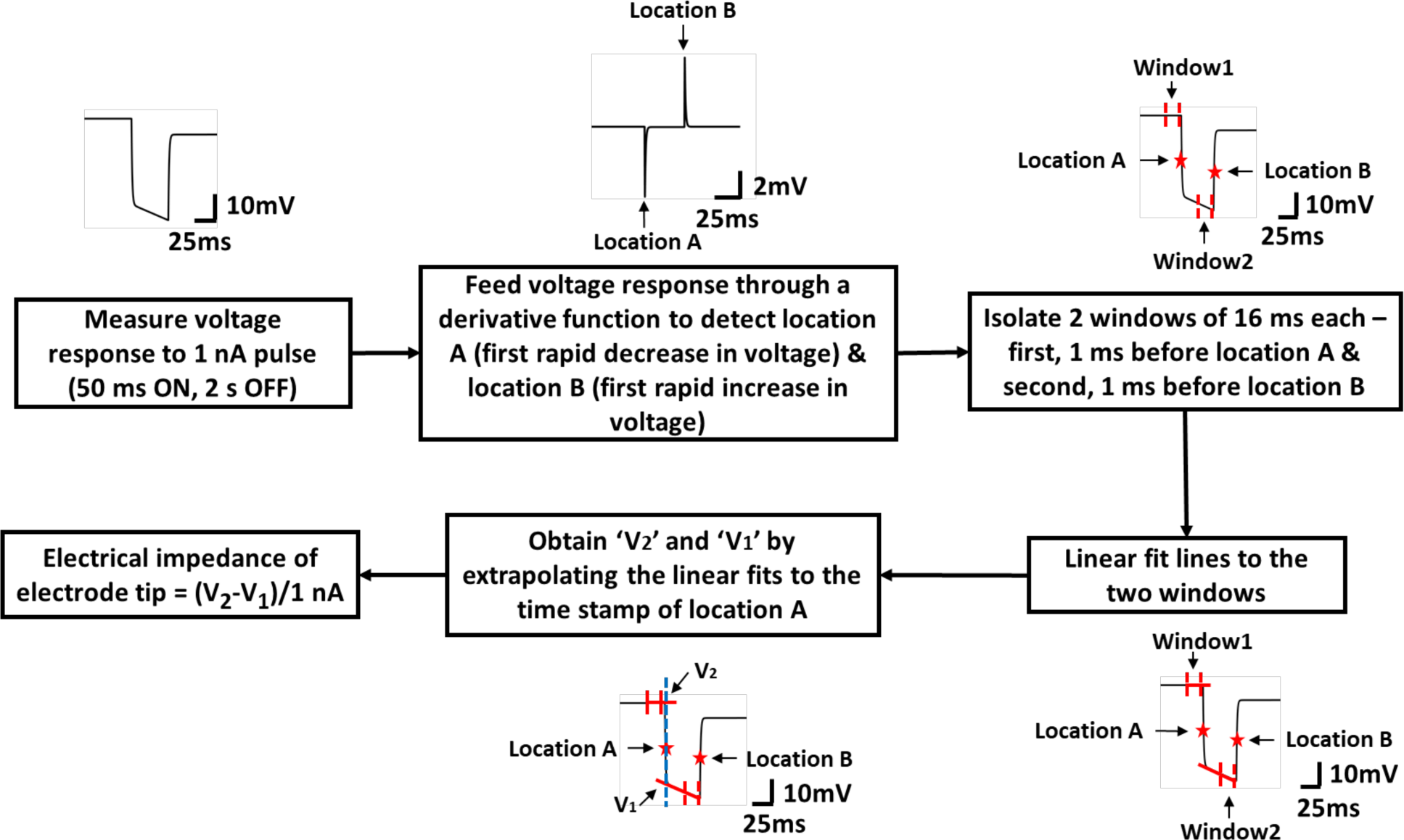
Modeling the electrical impedance of glass-polysilicon (GP) microelectrode. (A) Schematic representing the elements of the microelectrode-neuron interface. (B) Equivalent electrical circuit of the electrode-neuron interface. *R*_*seal*_ represents the degree of coupling between the electrode and neuronal membrane. (C) Expected (ideal) voltage responses of this electrode to application of a 1 nA current pulse (shown in inset) for 2 conditions: (i) Seal resistance (*R*_*seal*_) of 0.1 MΩ (simulating a condition of electrode away from neuron), (ii) Seal resistance (*R*_*seal*_) of 1 GΩ (simulating a condition of electrode inside neuron). (D) measured voltage response of a GP microelectrode to the application of a 1 nA pulse, matches well with the response predicted by the model for *R*_*seal*_ = 0.1 MΩ. (E) Sequence of steps involved in quantitative measurement of electrical impedance of the tip of the glass-polysilicon microelectrode.

The expected voltage responses of the electrode to a 1 nA pulse (100 msec ON, 2 sec OFF) were obtained from the model for 2 different values of seal resistance (*R*_*seal*_) (Fig. 3C). For a *R*_*seal*_ of 0.1 MΩ, which simulates a condition of electrode being at a distance from the cell, the response had an initial rapid decrease in voltage, followed by a slower decrease (Fig. 3C(i)). The rapid decrease with a time constant (τ_1_) of 0.5 msec corresponds to the RC combination at the miniaturized glass micropipette (*R*_*tip*_.*C*_*d*_). The subsequent slower decrease with a large time constant (τ_2_) corresponds to the RC combination at the polysilicon-electrolyte interface (*R*_*ct*_.*C*_*dl*_). Therefore, the electrical impedance of the tip of the electrode for this condition is proportional to the magnitude of voltage response after the first rapid decrease (at least 3^*^ *R*_*m*_.*C*_*m*_) for the initial steady state).

A *R*_*seal*_ of 1 GΩ simulates a condition of the electrode inside a cell. The corresponding voltage response had three distinct time constants as shown in Fig. 3C(ii). The first time constant corresponds to the RC combination at the miniaturized glass micropipette τ_a =_ *R*_*tip*_.*C*_*d*_, the second decrease with a time constant of 12 msec is due to the cell membrane τ_b_ = *R*_*m*_.*C*_*m*_ and a third slow decrease is due to polysilicon-electrolyte interface τ_c_ = *R*_*ct*_.*C*_*dl*_. Thus, the first and second decreases in the voltage response correspond to events that occur at the tip of the electrode due to interactions between the electrode and cell, while the third decrease corresponds to events at the polysilicon-electrolyte interface. For this condition, the electrical impedance of the tip of the electrode is proportional to the magnitude of voltage response after the second decrease (at least 3* *R*_*m*_.*C*_*m*_) for the RC network to reach steady state).

The voltage response predicted by the model for *R*_*seal*_ = 0.1 MΩ was validated experimentally by measuring the response of an electrode to a 1 nA current pulse (50 msec ON, 2 sec OFF) applied via the intracellular amplifier. The measured response closely followed the predicted response (*R*^2^ = 0.98 and normalized root-mean-square error, NRMSE = 0.05) (Fig. 3D). NRMSE was computed by dividing the RMSE by the range of the predicted response.

### Quantitative measurement of electrical impedance of the tip of the electrode

Based on the predicted voltage responses of the GP microelectrode shown in Fig. 3C, we developed an algorithm for quantitative measurement of electrical impedance of the tip of the electrode that is illustrated in the schematic of Fig. 3E. The duration of 1 nA current pulses is set to at least 50 msec to allow for the RC combinations of *R*_*tip*_.*C*_*d*_ and *R*_*m*_.*C*_*m*_ to reach steady state. The response to the current pulse is fed through a derivative function to capture the locations of the first decrease and first increase in voltage. The algorithm isolates two 16 msec windows from the voltage response: i) just prior to the first decrease and, ii) just prior to the first increase. Linear fits for these windows are obtained and extrapolated to the timestamp of the location of the first decrease to obtain V_1_ and V_2_ as shown in Fig. 3E. The difference in their corresponding voltage magnitudes is proportional to the electrical impedance of the tip of the electrode (Z_tip_ = (V_2_ - V_1_)/1 nA).

### Tracking electrical impedance of the tip of GP microelectrode as it approaches a neuron

To evaluate the efficacy of the electrical impedance of the tip of GP microelectrode as a feedback variable for the autonomous positioning algorithm, we used the model (Fig. 3B) to predict the electrical impedance of the GP microelectrode approaching a neuron. The increase in proximity of the electrode to a neuron was modeled as an increase in seal resistance from 0.1 MΩ to 1 GΩ. For each *R*_*seal*_, voltage output of the model in response to 1 nA input current pulses (50 msec ON, 2 sec OFF) was predicted and the electrical impedance of the electrode tip was calculated from the corresponding voltage response using the algorithm shown in Fig. 3E. The change in electrical impedance of the electrode tip (subtracted from the initial value of electrical impedance of the electrode tip for *R*_*seal*_ = 0.1 MΩ) plotted against *R*_*seal*_ is shown in Fig. 4A. For values of *R*_*seal*_ < 250 MΩ, the electrical impedance of the electrode tip increased monotonically with increase in *R*_*seal*_. The electrical impedance increased to 21 MΩ (close to the value of *R*_*m*_ in the model = 25 MΩ) at *R*_*seal*_ = 250 MΩ. For *R*_*seal*_ larger than 250 MΩ (up to 1 GΩ), only marginal increase in the electrical impedance was predicted. Predicted voltage responses of the model to 1 nA current pulses for 5 increasing values of *R*_*seal*_ are also shown in Fig. 4A.

**Fig.4:**
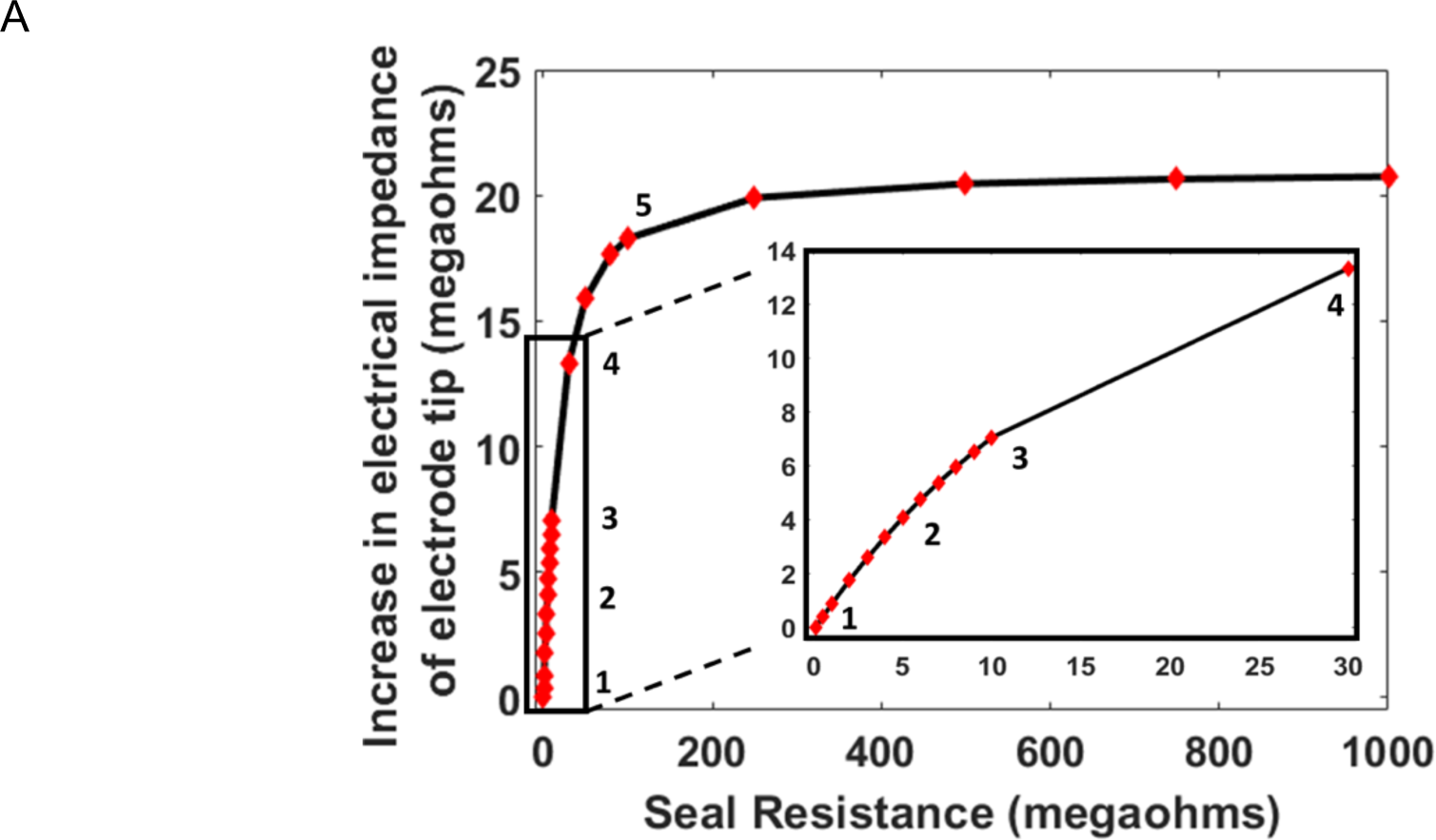
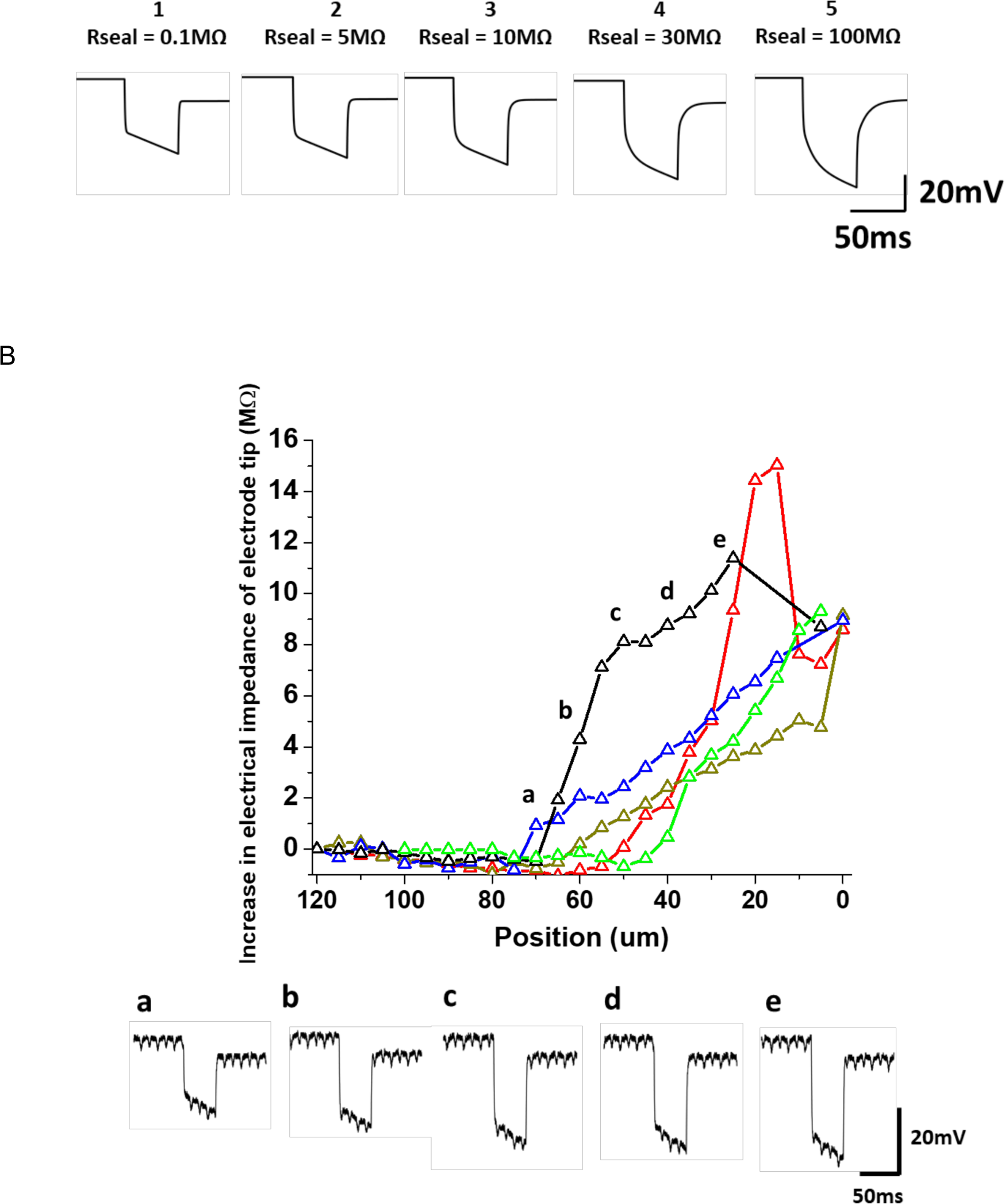
Electrical impedance of the tip of GP microelectrode as it approaches a neuron in the isolated abdominal ganglion. (A) The decreasing distance between electrode and neuronal membrane was simulated by increasing the seal resistance, *R*_*seal*_.) For each *R*_*seal*_, voltage response of the GP microelectrode to a 1 nA current pulse was predicted and the electrical impedance of the electrode tip was measured using the algorithm in Fig. 3E. The voltage responses predicted by the model for five increasing values of *R*_*seal*_ is shown below. (B) Experimental measurements of change in electrical impedance of the tip of the GP microelectrodes with increase in proximity to a neuron (n = 5 neurons). The increase in electrical impedance began 45-70 μm before successful cell penetration across the five electrodes. The recorded voltage responses of an electrode at 5 different positions of the electrode with respect to a neuron is shown in the lower trace.

Experimental measurements of electrical impedance of the tip of the GP microelectrodes (n = 5) as they approached abdominal ganglion neurons (5 cells) validated the impedance trend predicted by the model (Fig. 4B). Electrical impedance of the tip of the electrodes at each position was computed from the recorded voltage responses to 1 nA pulses (50 msec ON, 2 sec OFF) using the algorithm described in Fig. 3E. Position ‘0 μm’ indicates the location where there was successful penetration of neuron and intracellular signals were observed for the first time. In 2 out of 5 electrodes, we were unable to record electrical impedances at the tip of the GP microelectrode at ‘0 μm’ position. For all GP microelectrodes, electrical impedance of the tip increased to > 8 MΩ before successful penetration of cell membrane. The increase in electrical impedance began 45-70 μm before successful cell penetration across the five electrodes. The monotonic increase in electrical impedance of the electrode tip indicated that it can be reliably used to detect proximity to a cell. Recorded voltage responses at 5 different locations of an electrode with respect to a neuron are also shown in Fig. 4B.

### Closed loop control validation using GP microelectrode and conventional microdrive

As a first step in the validation, the closed loop controller was integrated with a GP microelectrode and a conventional hydraulic microdrive in the closed-loop scheme outlined in Fig. 8A. A typical trial using this integrated system is illustrated in Fig. 5A. When the controller was initiated, it operated in the ‘neuron search’ mode and moved the electrode in 5 μm steps while measuring electrical impedance of the tip and membrane potential after each step. As soon as the electrical impedance of the electrode tip increased above the set threshold (8 MΩ for this trial), the controller switched to the “penetration and/or tuning” mode and moved the electrode by 20 μm at a speed of 40 μm/s to enable penetration. Subsequently, the controller transitioned to the ‘Maintain’ mode as soon as good quality intracellular signals (RP < −35 mV and/or AP > 60 mV) were recorded. The performance of the controller in 2 additional neurons is demonstrated in Fig. 5B. In each trial, the controller successfully isolated and penetrated a neuron as well as obtained good quality electrical recordings. It should be noted that the controller moved the electrode by one step every 30 sec in the ‘neuron search’ mode. The ‘time’ axis in the plots have been truncated to better represent the increase in electrical impedance of the electrode tip over successive steps during the ‘neuron search’ mode. The actual time taken by the algorithm to move the electrode by 80 μm was 7.5 min.

**Fig.5:**
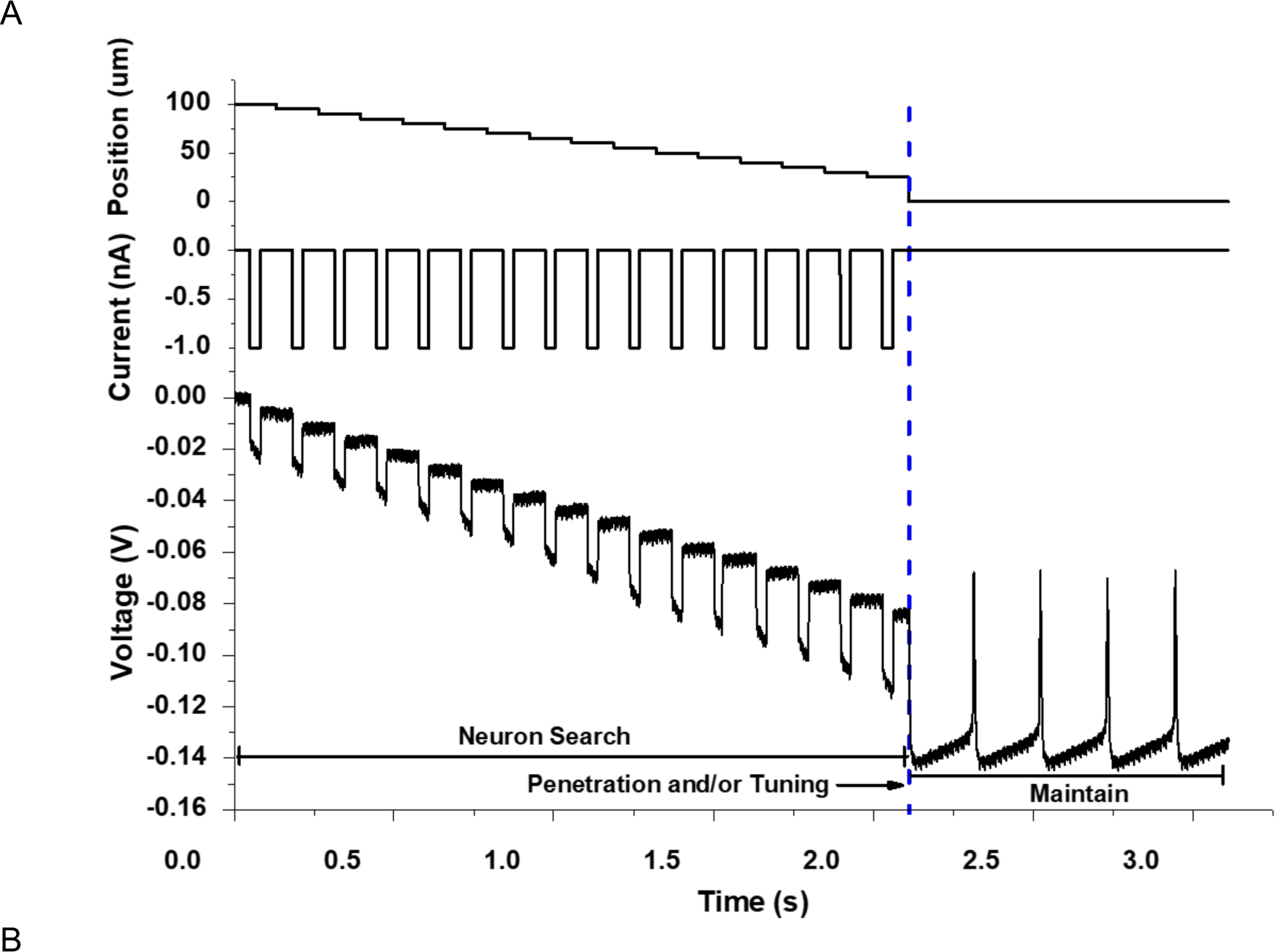
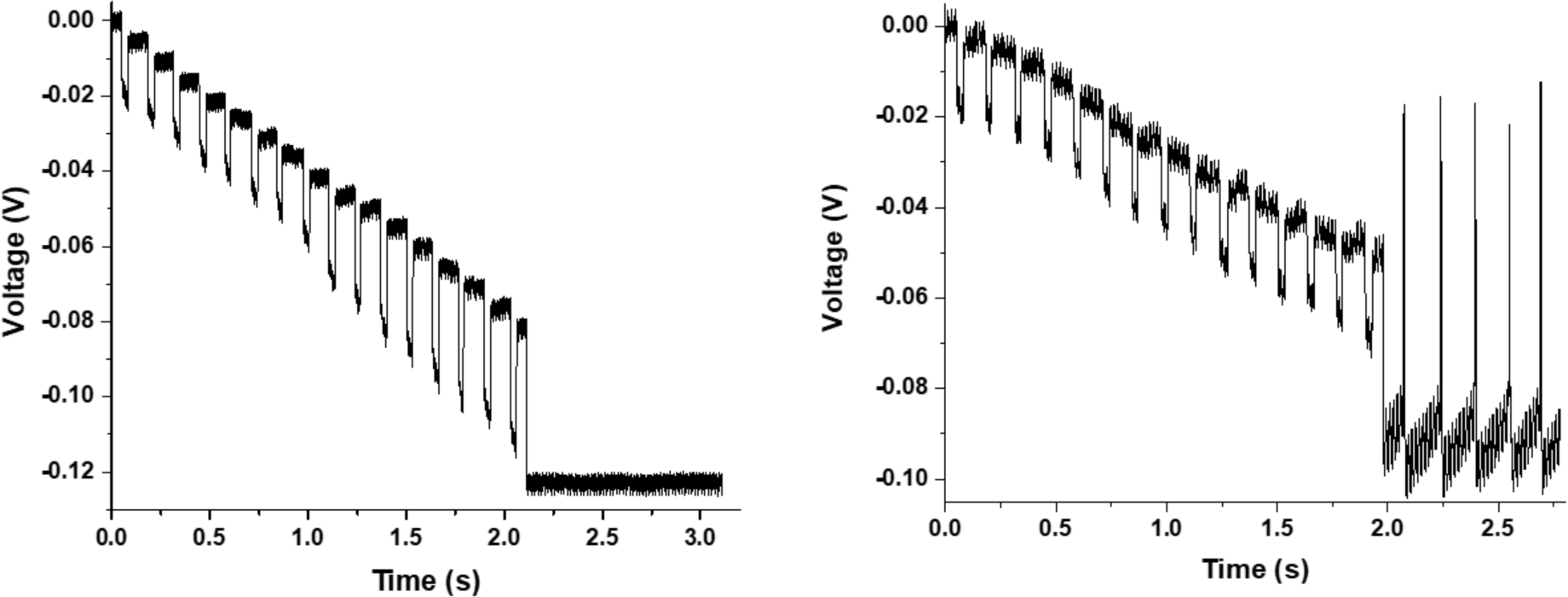
Validation of the closed-loop control after integration of controller with a <u>GP microelectrode</u> and a <u>conventional hydraulic microdrive</u>. (A) Illustration of a typical trial of the closed loop control - the controller moved the electrode using the microdrive in steps of 5 μm towards a neuron in the isolated abdominal ganglion, until the electrical impedance of the tip was above the threshold value of 8 MΩ, beyond which the controller switched to the ‘Penetration and/or Tuning’ mode to impale the neuron and obtained intracellular recordings. (B) Representative plots showing performance of the controller in 2 additional neurons with good quality resting potentials (RP) and/or action potentials (AP).

### Forces required to penetrate neurons in the abdominal ganglion of *Aplysia Californica*

To assess the ability of electrothermal microactuators to penetrate single neurons in the abdominal ganglion of *Aplysia Californica*, we measured the forces required to penetrate the neurons with a GP microelectrode at different electrode movement speeds. Fig. 6A shows simultaneous recordings of force and voltage during insertion and removal of a GP microelectrode from a neuron. Fig. 6A(i) shows the forces acting on the GP microelectrode during different stages of electrode interaction with the cell. During downward movement of electrode into the cell, forces were registered as negative values by the load cell due to compression. During upward movement of electrode out of the cell, forces were registered as positive values due to tension. There was minimal relaxation of neuronal membrane around the electrode (< 10 μN) after successful penetration of neurons over the duration of our force recordings. Forces required to penetrate the neuron was therefore, measured as the maximum decrease in the force curve. Only neurons from which good quality intracellular potentials were recorded upon penetration were included in the experiment. Penetration forces were measured from n=5 neurons (3 animals) for 7 different electrode movement speeds. For 1 of the 5 neurons, we were able to measure penetration forces only for 3 different speeds. Only one trial was performed at a given speed to reduce damage to the neuron due to repeated penetration. We did not observe any significant trend in the measured forces as a function of penetration speed in a given neuron (Fig. 6B). Therefore, force measurements from 3 different speeds were pooled and box-whisker plots of penetration forces for the five neurons are shown in Fig. 6C. The median forces required for the penetration of five neurons tested were 32 μN, 60 μN, 143 μN, 75 μN and 152 μN indicating a possible dependency on cell-type. Tukey test results for pairwise comparisons of mean penetration forces for the five neurons showed that the forces were significantly different for 8 out of 10 pairs of neurons (*p*< 0.05) (Fig. 6D). The force generated by the electrothermal microactuators during one stroke has been estimated from our previous studies to be 111 μN per electrothermal strip [25]. The actuators used in this study had at least 2 electrothermal strips and were capable of generating a force of 222 μN, which was significantly higher than the forces required to penetrate abdominal ganglion neurons.

**Fig.6:**
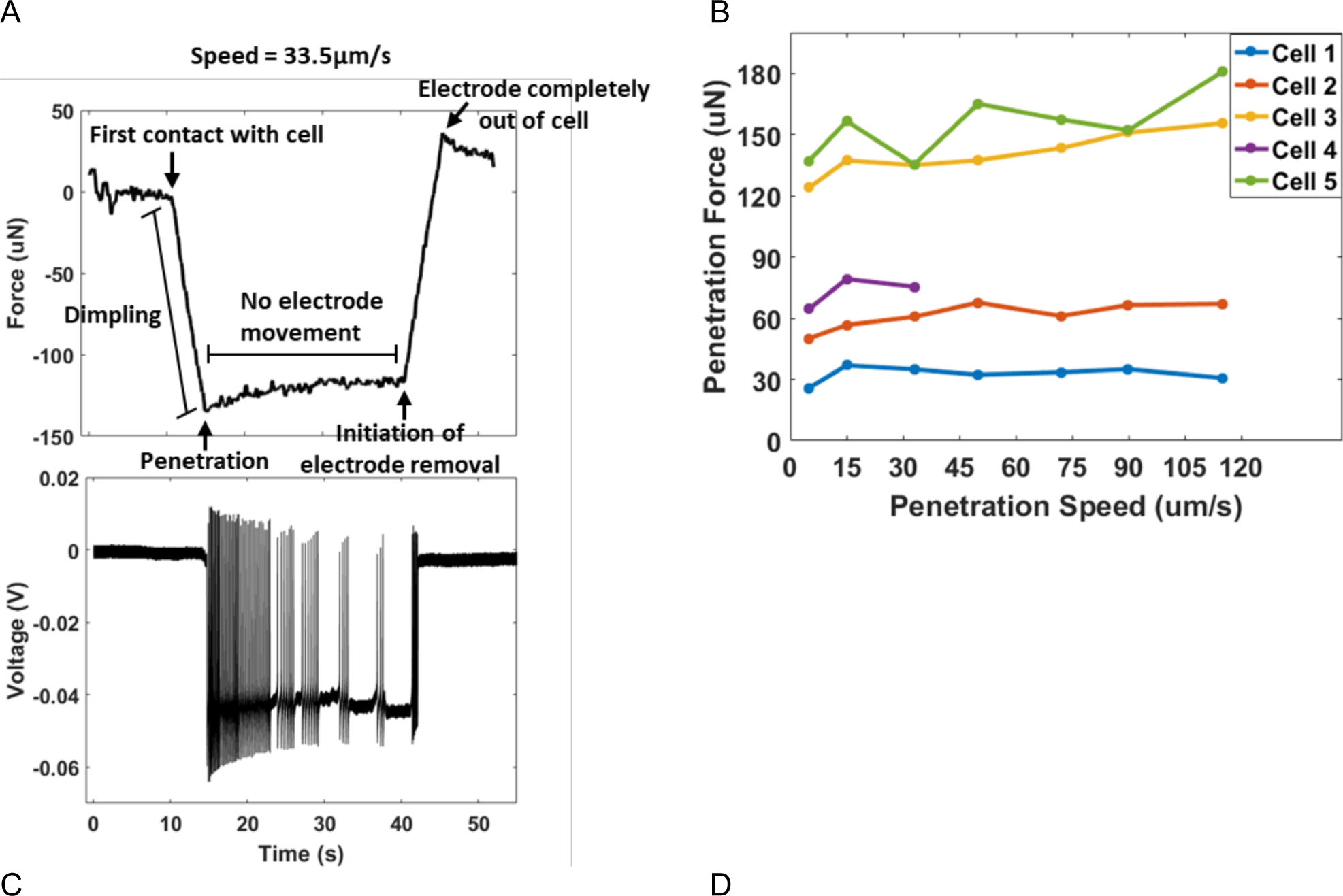
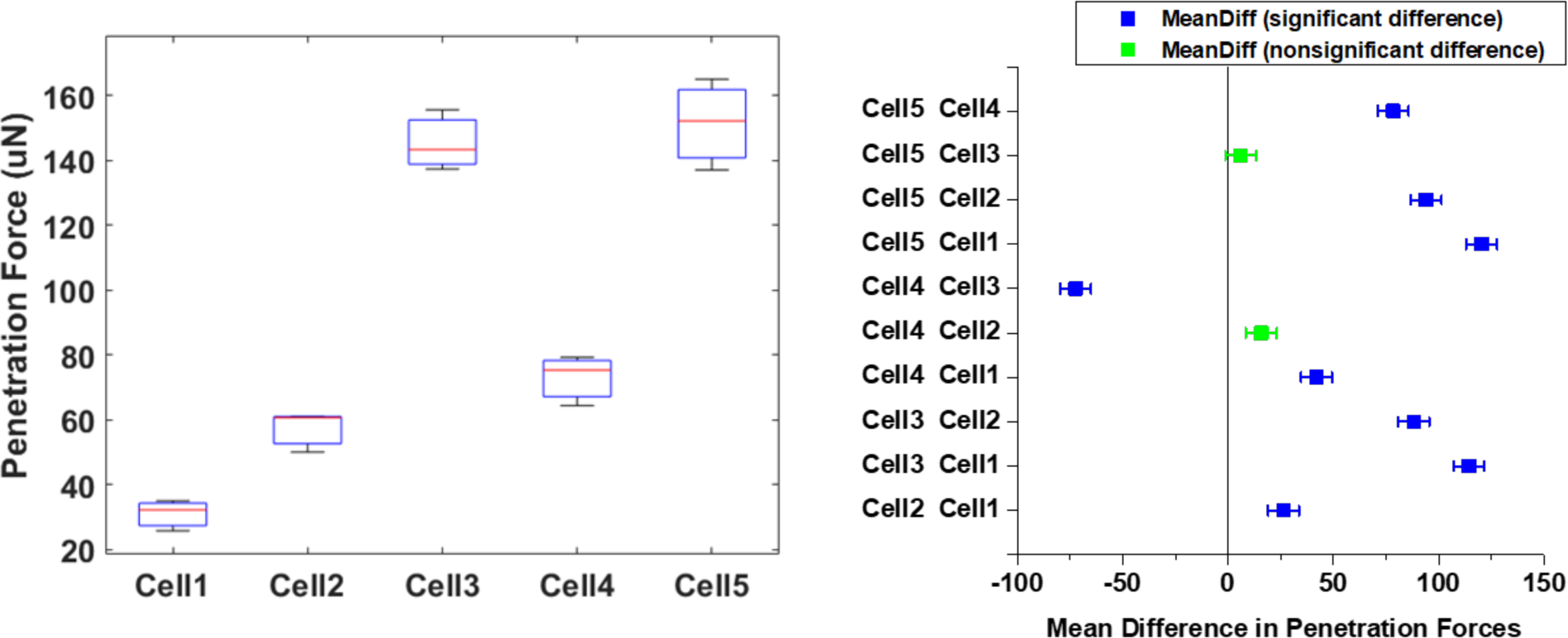
Cell penetration forces in the neurons of isolated abdominal ganglion show significant variation among neurons with median forces ranging from 32-152 μN. (A) Simultaneous force and voltage recording during insertion and subsequent removal of a GP microelectrode from cell. (B) Penetration forces showing minimal variability across different electrode movement speeds in n=5 neurons. (C) Box-whisker plots of penetration forces measured for 5 different neurons in the abdominal ganglion. (D) Tukey test results for pairwise comparisons of mean penetration forces for all neurons tested. The forces were significantly different for 8 out of 10 pairs of neurons (*p*< 0.05).

### Validation of closed loop control integrated with the MEMS-based microactuators

In the final set of closed loop control experiments, the closed-loop controller was integrated with the 2 MEMS-based sub-systems - the GP microelectrode and electrothermal microactuators. In the ‘neuron search’ mode, the controller moved the electrode towards a cell in 6.5 μm steps using the microactuators. After an increase in electrical impedance of the tip of the electrode above threshold (6 MΩ), the controller autonomously penetrated the cell with a continuous 26 μm movement (speed = 40 μm/s), before switching to the ‘maintain’ mode during which good quality intracellular recordings were observed as shown in Fig. 7A. The controller, integrated with the MEMS-based sub-components, repeatably searched, penetrated and recorded high fidelity intracellular potentials from 3 different abdominal ganglion neurons as shown in Fig. 7. As noted before, the ‘time’ axis in the plots have been truncated to show the increase in electrical impedance of the tip of the GP microelectrodes across steps. The actual time taken by the controller to move the electrode by 80 μm using the microactuators was 6 min.

**Fig.7:**
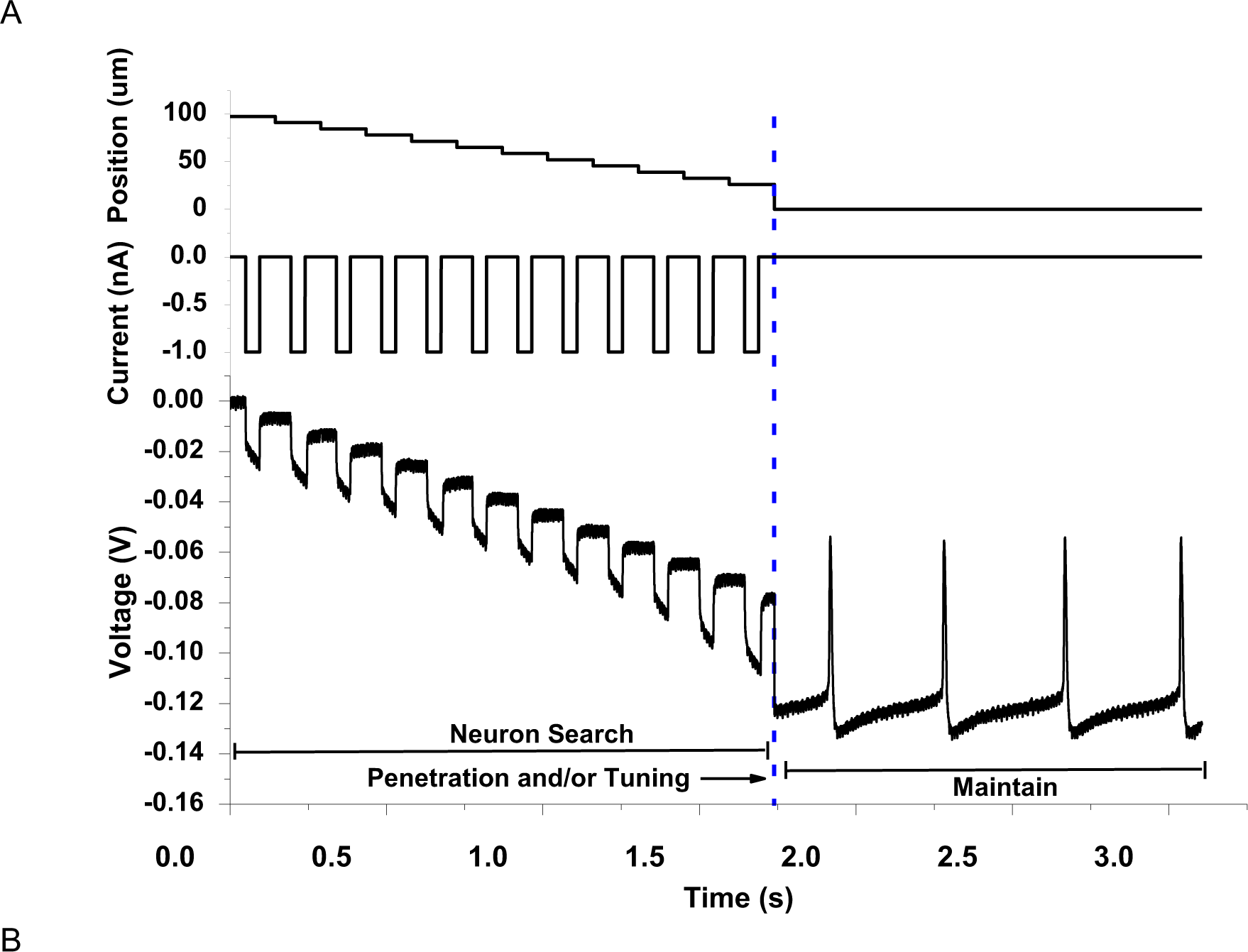
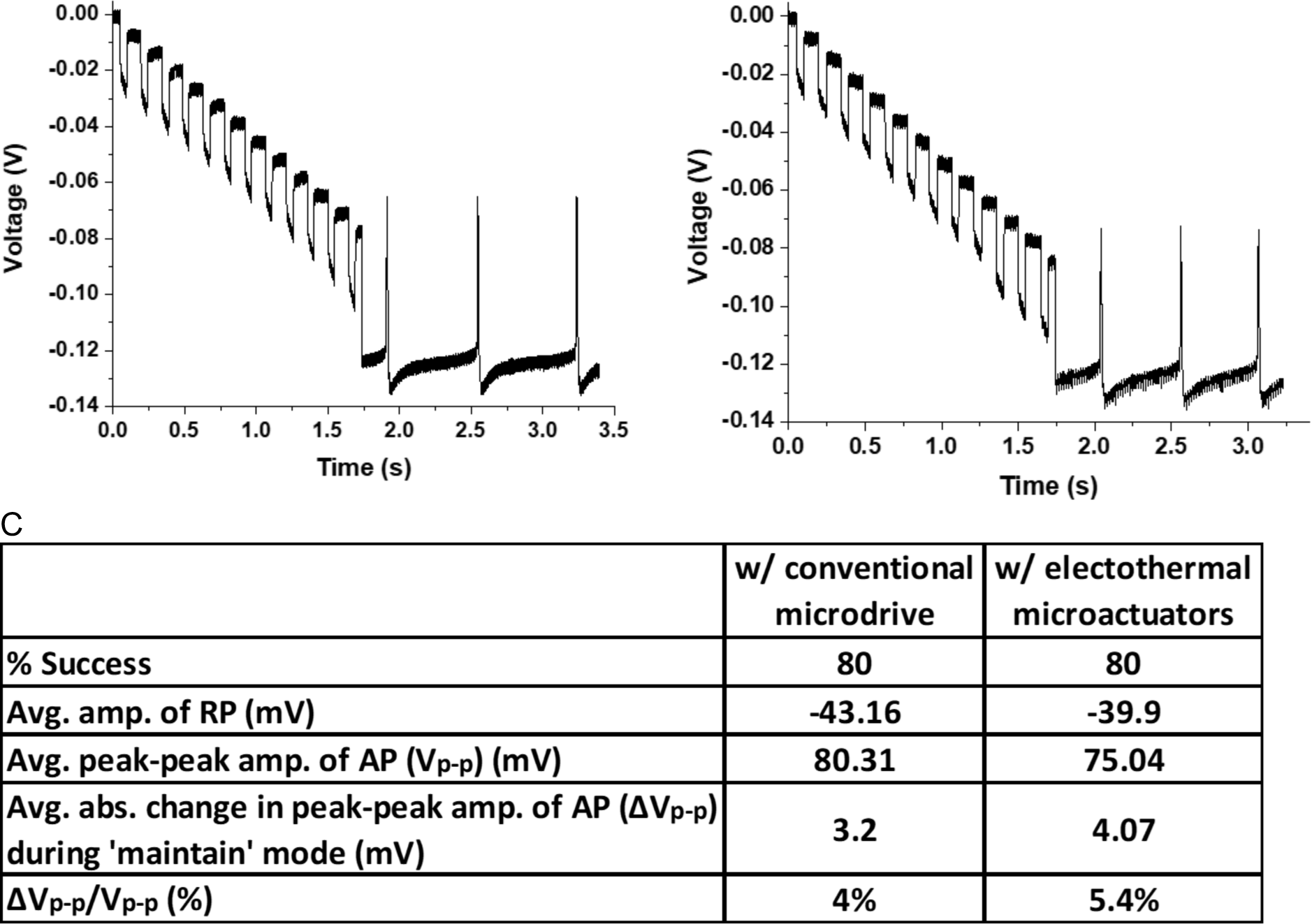
Validation of closed loop controller integrated with the two MEMS-based sub-components - <u>GP microelectrode</u> and <u>electrothermal microactuators</u>. (A) A typical trial of the closed-loop controller - the controller moved the electrode using the electrothermal microactuators in steps of 6.5 μm towards a neuron until the electrical impedance of the electrode tip was below the threshold value of 6 MΩ, after which the controller switched to the ‘penetration and/or tuning’ mode to autonomously penetrate the neuron and record intracellular potentials. (B) Two additional trials demonstrating ability of the controller and microactuators to successfully locate and penetrate 2 separate neurons. (C) Comparison of performance of the controller with the conventional microdrive (5 neurons) vs. electrothermal microactuators (5 neurons) showed no significant difference in RP, AP, ΔVp-p between the two systems (*p*>0.1 in all cases).

**Fig.8:**
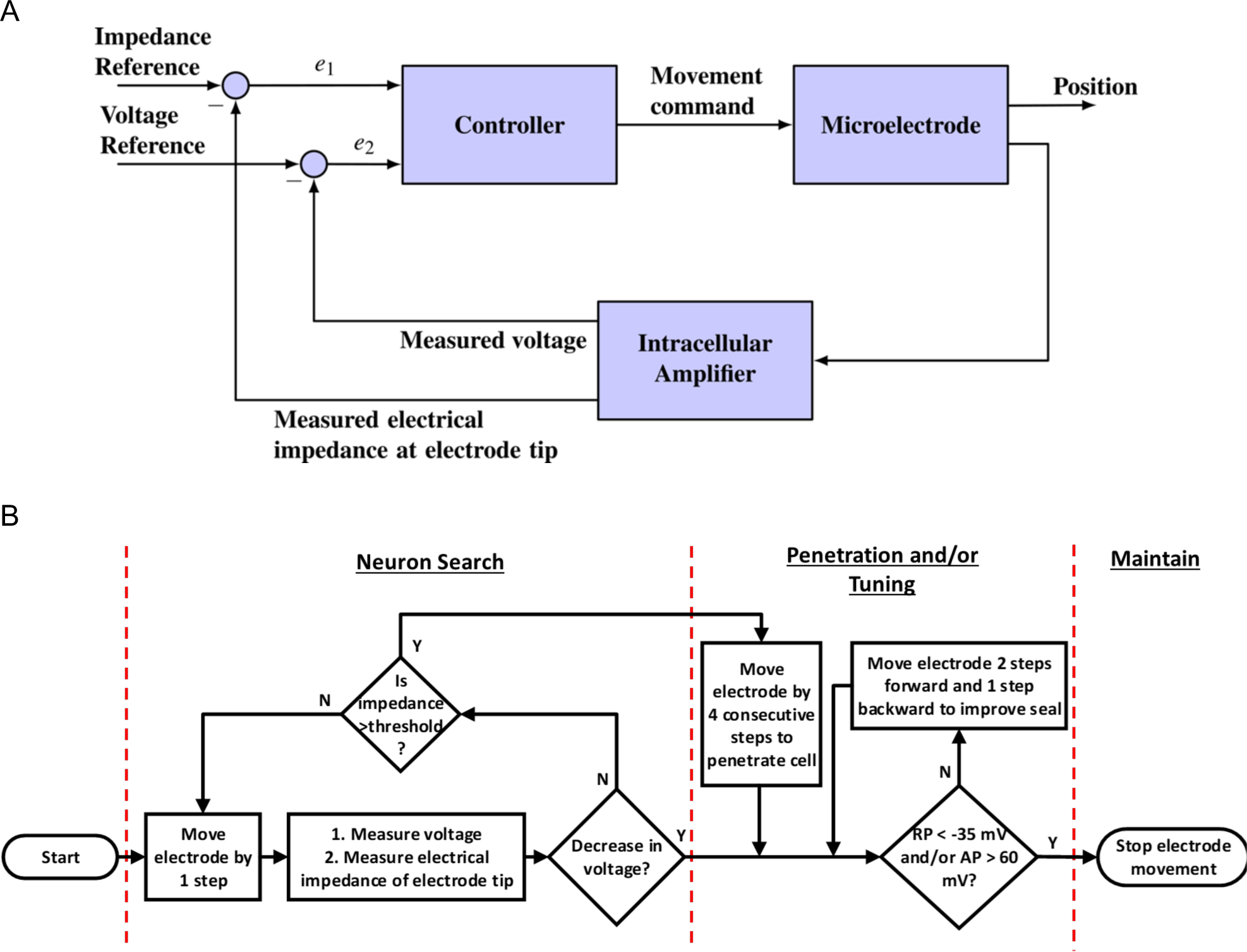
Closed loop control algorithm for autonomous movement and positioning of glass-polysilicon microelectrode inside single neurons. (A) Block diagram of the ON-OFF closed-loop control scheme with two feedback variables-electrical impedance of the tip of the electrode at DC and voltage recorded at tip of the electrode (B) Flowchart showing the steps involved in the three modes of the control algorithm: i. Neuron Search mode, ii. Penetration and/or Tuning mode and iii. Maintain mode (RP: Resting Potential, AP: Action Potential)

### Comparison of controller performance with conventional microdrive vs electrothermal microactuators

Fig. 7C compares the performance of the controller integrated with the conventional hydraulic microdrive system (5 neurons) with that of the controller integrated with the electrothermal microactuator system (5 neurons). In all neurons, the recordings were terminated 15 min after the initiation of the ‘maintain’ mode to perform more trials. With both motors, the controller was successful in 4 out of 5 neurons (80% yield). The quality of recordings obtained after autonomous penetration of neurons with the two different systems were comparable. There was no statistically significant difference between the means of recording (RP and peak peak AP) amplitudes obtained with the two different systems (RP: *p*>0.2, AP: *p*>0.1). Further, the maximum absolute change in peak-peak amplitudes of AP over the duration of 15 min in ‘maintain’ mode were not significantly different for the two systems (*p*>0.2).

## Discussion

This study demonstrates a fully autonomous, MEMS-based intracellular recording system that integrates a polysilicon-based microelectrode, microscale electrothermal actuators and closed-loop control for intracellular recordings from single neurons. The system has been validated in the abdominal ganglion neurons of *Aplysia Californica*, and will be further validated in vivo rodent experiments in the future. The form factor of the current system will allow it to be mounted on the head of a rodent in long-term experiments meeting the long-standing need for a head-mountable, autonomous intra-cellular recording system.

We first demonstrated the ability of the GP microelectrode to record high fidelity intracellular signals from abdominal ganglion neurons (Figs. 2A, 2B) as well as the motor cortex of a rat in vivo (Fig. 2C). The short in vivo recording, spanning 7 sec, was possibly recorded from a glial cell rather than a motor neuron based on the marginally hyperpolarized resting potential (−74 mV) and absence of action potentials [29], [30]. Further rigorous evaluation of the ability of this GP microelectrode to reliably record from neurons in vivo is required.

### Forces required to penetrate the Aplysia Californica neurons

In order to evaluate the force demands on the electrothermal microactuators, the forces required to penetrate neurons of *Aplysia Californica* using GP microelectrodes were measured. Neuron-penetration forces ranged 32 μN to 152 μN in this study (Fig. 6C). Previous studies report a large range of penetration forces across various live cell types ranging from < 1 nN to 30 nN [31]–[36]. These studies typically used rigid AFM probes with a variety of tip shapes (pyramidal, conical, cylindrical), lengths of 6-10 μm and tip diameters of 30-300 nm to penetrate plated cells with heights of 5-10 μm. The current study used GP microelectrodes 5-6 mm long having conical-frustum-shaped tips of ∼500-600 nm O.D. Angle et al [31] and Obataya et al [32] measured higher penetration forces with increase in tip dimensions of the probes. Neurons of *Aplysia Californica* are large (250-600 μm diameter) with heights comparable to their diameters as they were not plated. Yokakowa et al [38] and Guillaume-Gentil et al [34] reported increase in penetration forces with increase in height/size of cells for the same probe. Therefore, we speculate that the large magnitude of forces observed here may have been the result of larger tip sizes of the electrode, larger diameters and heights of neurons used in this study. We used four different electrodes to obtain penetration forces from five different neurons 250-500 μm in diameter (3 animals). The large variation in the measured forces across neurons can be attributed to a variety of factors, such as electrode-to-electrode variability, location of electrode with respect to cell soma (center vs. corner), size of cell, health of cell, location of cell in ganglion (how tethered it is) and effect of protease (for digestion of connective tissue sheath covering the ganglion) on neuronal membrane stiffness. Angle et al. [31] also observed large variations in measured forces (∼3.2–32 nN) using rigid AFM probes for a given cell type, electrode type and experimental conditions. Similar penetration forces across a range of penetration speeds 5 – 125 μm/s (Fig. 6B) suggest that the neuronal membrane breaks down above a certain stress, irrespective of the strain rate within the penetration speeds tested here. Interestingly, Kawamura and colleagues noted that the approach velocity (1-1000 μm/s) of AFM nanoneedle probes had no significant effect on the success rate of penetration of HeLa cells [39]. At penetration speeds of 5 – 125 μm/s, force data in the current study suggests that the *Aplysia Californica* neurons indented by approximately half their diameter (150-300 μm) before penetration. Yokokowa et al. [38] reported a positive correlation between cell height and indentation depth prior to cell penetration for the same probe. The lack of any significant relaxation in forces after neuronal penetration indicate that the neuronal membrane remains dimpled over the duration of our force recordings lasting up to 10 mins, possibly due to the tight seal between glass and cell membrane. Prolonged dimpling of cell may have implications on the functional state of cell, such as activation of membrane stretch receptors.

### Electrical impedance of the tip of GP microelectrode

Simulation results (Fig. 4A) suggest that the increase in electrical impedance of the tip of the electrode is an effect of increasingly tight coupling between the electrode and neuron, due to which larger fraction of the current being applied to the electrode (to estimate electrical impedance of the tip of the electrode) flows through the cell (via the membrane) and smaller fraction flows through the extracellular space. The increase in electrical impedance of the electrode tip was close to the membrane resistance (*R*_*m*_) of the cell for values of *R*_*seal*_ > 250 MΩ, indicating maximum electrode-neuron coupling and most of the current was flowing through the cell. Increase in electrical impedance of the tip of conventional patch and sharp glass micropipettes has been reliably used to detect neuron proximity in vivo [8], [10]–[12]. In the current study, experimental measurements of electrical impedance of the tips of GP microelectrodes increased monotonically to > 8 MΩ over a distance of 45-70 μm before successful penetration of the neurons (Fig. 4B). In contrast, Ota et al. [11] observed similar increases in electrical impedance of the tips of sharp micropipettes over smaller distances of 10-20 μm before penetration of mouse cortical neurons in vivo. The large size of *Aplysia Californica* neurons (250-500 μm in diameter) and larger indentation depth (150-300 μm) before penetration likely caused the electrical impedance to increase over larger distances before penetration than prior studies. Here, five different electrodes were used to record electrical impedances from five different cells. The large variation in electrical impedances across the 5 different GP microelectrodes in Fig. 4B and the distance over which the impedance increases (45-70 μm) could be due to variabilities in size of neurons and location of electrode with respect to cell soma (center vs corner). We measured cell input resistances of ∼25-28 MΩ in *Aplysia Californica* neurons when full-blown action potentials were recorded (data not included). However, the increase in electrical impedances at position ‘0 μm’ in Fig. 4B were much smaller (8-10 MΩ) possibly because the electrode tips were not completely inside the cells. This was corroborated by the smaller-amplitude action potentials (10-20 mV) and depolarized resting potentials (−10 to −15 mV) recorded at that position. Finally, the increase in electrical impedance as a function of distance from the neuron was not monotonic for a short segment in two of the 5 GP microelectrodes in Fig. 4B. One explanation for this is that the GP microelectrodes in the above 2 cases might have been slightly off-center with respect to the neuron, which may have resulted in a change in contact area between the electrode and neuronal membrane with successive steps.

Currently, the controller waits for 30 sec between successive electrode movements to allow for the charge buildup at the polysilicon-KAc interface upon application of 5 consecutive pulses of 1 nA (50 msec ON, 2 sec OFF) for measurement of electrical impedance of the tip of the microelectrode to discharge completely. Therefore, the time taken to move the electrode by 100 μm would be ∼7.5 min (with the electrothermal microactuators) or 10 min (with the conventional motorized micromanipulator). The wait-time between steps can be potentially minimized by using biphasic, charge-balanced current pulses for measurement of electrical impedance of the tip of the microelectrode. Further, our data also indicates that the 1 nA currents used for impedance measurements did not alter the interface in any measurable way. Electrochemical impedance spectra of the GP microelectrode before and after continuous application of 1500 pulses (50 msec ON, 5 sec OFF) of 1 nA did not show any significant changes as assessed by their NRMSE values. Visual inspection of the GP microelectrode under the microscope showed no bubbles, indicating that the currents applied were well within the electrochemical window of KAc and did not result in electrolysis in the miniaturized glass micropipette. Further, high fidelity recordings as well as minimum change in the peak-peak amplitudes of AP of the recorded signals during ‘maintain’ mode of the controller in 8 neurons (Fig. 7C) indicate that the charging and discharging events had no adverse effects on the neurons being recorded from. The long-term stability of the interface has to be carefully evaluated in future in vivo studies.

### Comparison of the closed-loop controller with conventional microdrives versus one integrated with the electrothermal microactuator

Both types of closed-loop control systems were successful in 4 out of 5 neurons (Fig. 7C). The failed trial with the conventional microdrive had a GP microelectrode tip clogged with biological debris, which obscured the electrical impedance feedback. In the failed trial with the electrothermal microactuators, the GP microelectrode did not penetrate cell at the end of the ‘penetration’ mode even though the hallmark increase in electrical impedance of the tip of the electrode was observed. This could have been a result of the electrode contacting the corner of the cell at a more grazing angle resulting in unsuccessful penetration. The quality and stability of intracellular recordings (RPs and APs) obtained with the closed-loop controller using the electrothermal microactuators were equivalent to those obtained with the conventional microdrive (Figs. 5 & 7), indicating that the functional impact on the neurons due to successful penetration and stable maintenance of the GP microelectrode inside the neurons was comparable using both systems. A change in AP amplitudes when electrode is kept stationary within cell can be attributed to several reasons: 1) membrane slowly resealing around the electrode, 2) dysfunction induced by penetration and maintenance of electrode inside neuron and 3) possibility of drifts in the positioning systems or the neurons in the media. Similar change in AP amplitudes during ‘maintain’ mode with the two different closed-loop systems indicate that the electrothermal microactuators and conventional microdrives have comparable positioning drift and holding force. All the trials with the actuators showed an increase in the AP amplitudes with time, suggesting that resealing of the neuronal membrane around the GP microelectrode may have been the main contributor. We have observed similar changes in the AP amplitudes (2-5 mV) from *Aplysia Californica* neurons with conventional glass micropipettes.

This miniaturized, robotic system offers several significant advantages for potential use in vivo. The overall dimensions of the system (electrodes + positioning components) before packaging are 3 mm x 6.3 mm x 1 mm and weight is ∼35 mg, which makes it head-mountable for recordings from unrestrained, behaving animals such as rodents and non-human primates. Using conventional wire-bonding approaches for packaging, the system used in this study had a slightly large footprint (1.78 cm x 1.46 cm x 0.5 cm) for 3 independently movable GP microelectrodes. Using flip-chip based packaging methods reported earlier, the size can be further reduced to chip-scale [40], [41] and help realize higher channel counts (with tens and hundreds of GP microelectrodes) and throughput in future designs. Our current design accommodates 3 intracellular electrodes per chip, spaced approximately 800 μm apart. In future designs, the spacing between the electrodes can be further reduced to approximately 400 μm by using optimal 3D chip-stacking strategies. Our previous characterization study [25] showed that the actuators are robust and have consistent stepping over 4 million cycles of operation in bench-top experiments. The electrothermal actuators also generate enough force to penetrate and navigate the electrode through brain tissue [25], [26], [42].The use of electrothermal microactuators allows for robust and reliable microscale movement of individual electrodes within brain tissue also. We have already demonstrated the reliability of these electrothermal microactuators in long-term rodent experiments where we tested the ability of these microactuators to maintain a stable extra-cellular interface with single neurons in the in vivo brain over periods lasting as long as 13 weeks [42]. However, the ability of these electrothermal microactuators to isolate and precisely penetrate neurons in vivo has to be validated carefully in future experiments.

Further, movable intracellular probes can be repositioned to seek new neurons in a given region in the event of loss of signal, even after implantation. Also, electrothermal microactuators can help improve the stability of intracellular recordings by incorporating control strategies such as those reported by Fee [15], where the intracellular electrode was moved along with the neuron to compensate for brain micromotion. Integration of a microscale, head-mountable positioning system such as the one reported here with the closed-loop automation algorithm for neuron isolation and penetration makes the system plug-and-play and thereby usable by several neurophysiology labs. This system can also be used for other applications such as current injection, dye labeling and targeted delivery by loading the miniaturized glass micropipettes with dyes or pharmaceutical agents.

In the current design, the GP microelectrode has a length of 5-6 mm from the edge of the chip. The electrode can be extended further by 2.5-3 mm using the electrothermal microactuators. Therefore, the electrode can be used to record intracellular activity from cortical as well as deep brain areas in rodents in vivo. There is a 2 mm overlap between the polysilicon electrodes and miniaturized glass micropipettes, due to which the full translation range of the microactuators is reduced to < 3 mm. This can be improved by increasing the length of our polysilicon microelectrodes in future designs. A significant challenge that remains to be addressed is preventing clogging of tips of the miniaturized glass micropipettes of the GP microelectrodes with debris. With the current design, the miniaturized micropipette has to be replaced in the event of tip clogging. We are investigating alternative designs of polysilicon based intracellular probes. The electrothermal microactuators have a displacement resolution of 6.5 μm, which is slightly larger than the typical step sizes used by neuroscientists when seeking neurons for intracellular recording in vivo (2-3 μm). The larger displacement resolution can potentially reduce the number of neurons we can sample/ record from. In order to increase our probability of penetrating neurons using the microactuators, the control algorithm can be modified to use extracellular voltage recording as an additional feedback variable, as it can detect neuron proximity before an electrode makes physical contact with a neuron (which causes an increase in electrode electrical impedance of the tip of the electrode). Finer displacement resolution can be readily achieved in future designs by reducing the spacing between the teeth along the sides of the polysilicon electrode and by using appropriate transmission gears between the electrothermal microactuators and polysilicon microelectrodes.

## Methods

### Fabrication of the electrothermal microactuators and microelectrodes

The electrothermal microactuators and microelectrodes were fabricated by surface micromachining polysilicon using the SUMMiTV™ (Sandia Ultra-planar Multi-level MEMS Technology V) process at the Sandia National Laboratories, Albuquerque, NM. The fabrication process is explained in our earlier study [26]. Briefly, SUMMiTV™ is a five-layer, surface micromachining process that uses silicon substrate topped with an insulating layer of silicon dioxide and silicon nitride as the starting material. Five layers of polysilicon deposited over this base layer make up the mechanical/electrical layers and each layer is separated from the next by sacrificial silicon dioxide layers. The mechanical components were defined in all the five layers, while the microelectrodes were defined in polysilicon layers 2 and 3. After all the layers were micromachined, the devices were released in HF to remove the sacrificial oxide layers (wet-oxide etch). Anchor points established electrical and mechanical connections between the different polysilicon layers after the oxide layers were removed. Spring-type leads made electrical connections between the moving microelectrodes and stationary bond-pads. The polysilicon microelectrodes were highly doped (10^21^ /cm^3^) in situ with phosphorus (n-type), to make them highly electrically conductive. The final part after dicing containing three polysilicon microelectrodes and associated electrothermal microactuators reported in this study have dimensions of 3 mm x 7 mm. Each of the three polysilicon microelectrodes (50 μm wide, 4 μm thick and 5 mm long) can be moved bi-directionally over 6 mm (forward and backward) with a displacement resolution of 6.5 μm, by means of the electrothermal microactuators.

### Packaging of MEMS device

A 40-pin chip carrier (Spectrum Semiconductor Materials Inc., San Jose, CA) was diced on one side, in order to create an opening through which the microelectrodes can extend off the MEMS chip after it has been bonded to the carrier. The MEMS chip was wire-bonded to the chip carrier using gold wires. A glass covering was glued to the top of the chip carrier to protect the device and wire bonding. Then, the chip carrier was bonded to a custom designed PCB using surface-mount technology. A male omnetics connector (Part # A79040-001, Omnetics Connector Corporation, MN) was also bonded to the PCB, and it would connect to a female omnetics connector (Part # A79045-001) integrated with a custom-made connector board to interface with the intracellular amplifier and data acquisition system, in order to enable computer control of actuation and recording. The overall dimensions of the packaged device (Fig. 1D) are 1.78 cm x 1.46 cm x 0.5 cm and the weight of the device is ∼1.9 grams. After packaging, the microelectrodes were extended off the edge of the chip to a length of 2 mm.

### Fabrication of miniaturized glass micropipettes

Borosilicate capillary glass tubes (A-M Systems, Carlsborg, WA) with an outer diameter of 1.5 mm and inner diameter of 0.86 mm were cut into 3-inch pieces. The glass pieces were cleaned by soaking in 60% nitric acid overnight. Then, they were rinsed in four changes of deionized water and methanol and dried at 60°C in an oven. Glass micropipettes were pulled using the horizontal Flaming/ Brown Micropipette P-87 puller (Sutter Instruments, Novato, CA). The pulling parameters were set such that the micropipettes had a final tip resistance of 10-20 MΩ. The glass micropipettes were surface-modified through silanization with *N*-(2-aminoethyl)- 3-aminopropyltrimethoxysilane (EDA) (Sigma Aldrich catalog # 440302) in order to reduce stiction forces between polysilicon and the wall of the miniaturized glass micropipette during their integration. Briefly, the pulled pipettes were placed in a closed petri-dish and warmed in an oven at 400°F for an hour. Then, a vial containing 4 µl of EDA was introduced into the closed petri-dish and pipettes were exposed to the silane vapor for 6 minutes. Following that, the vial was removed, and the oven door was kept open for about 30 sec to release any excess silane vapor. Then, the pipettes were baked for another 30 min at 400°F. After silanization, the electrodes were filled with 2M potassium acetate (KAc) solution that was degassed in a vacuum chamber for 6 hours. Air pressure was applied by connecting the broad end of the pipettes to an air power dispenser system, Ultimus I from Nordson Engineering Fluid Dispensing (EFD, part # 7017041) in order to fill the very tip of the pipettes. After filling, the tips of glass micropipettes (length of 4-5 mm and diameter of about 80-100 µm at the broad end) were carefully broken.

### Integration of polysilicon microelectrode with miniaturized glass micropipettes

The miniaturized glass micropipette was carefully mounted at the end of a vacuum pick up syringe, which was mounted on a 3-axis manual micromanipulator (ALA Instruments). The MEMS device with the extended polysilicon microelectrode was placed on a custom-built stage with microscale control in x, y and z directions. The miniaturized pipette and polysilicon electrode were aligned using cameras in x, y and z directions and the miniaturized pipette was carefully inserted at the end of the polysilicon electrode using the stage controls. After insertion, the broad end of the miniaturized pipette was sealed with a two-part silicone gel (3-4680, Dow Corning, Midland, MI) mixed in a ratio of 1:1 to prevent rise of electrolyte in the pipette into the MEMS device.

### Closed loop control algorithm

The closed-loop control scheme for autonomous navigation and positioning of microelectrodes inside single neurons in the isolated abdominal ganglion of *Aplysia Californica* is illustrated in Fig. 8A. This is an ON-OFF controller in which electrical impedance of the electrode tip at DC and voltage recorded at the tip of the electrode serve as feedback variables, as the electrode is advanced in micrometer steps towards a cell.

#### Modes of control algorithm

The control algorithm operates in three different modes, as illustrated in Fig. 8B:

##### 1. Neuron search

The controller is in the ‘neuron search’ mode until the electrical impedance of the tip of the electrode is above a set threshold value or until there is a decrease in recorded voltage (indicating cell penetration). In this mode, the controller moves the electrode in 6.5 μm steps. There is a 30 sec wait time between movements to measure the electrical impedance of the tip of the electrode, resulting in a movement speed of 13 μm/min.

##### 2. Penetration and/or tuning

If there is a spontaneous cell penetration at the end of the ‘neuron search’ mode and the electrical impedance of the tip of the electrode is above a pre-determined threshold (indicative of contact with cell membrane from prior experiments), the controller moves the electrode by 4 consecutive steps of 6.5 μm at a speed of 40 μm/s to allow cell penetration. After penetration, the controller evaluates the quality of signals recorded in a 10 sec segment. If the amplitude of resting potential is >-35 mV and/or the peak-to-peak amplitude of action potentials is < 60 mV, the controller moves the electrode two steps forward and one step backward to improve quality of recordings.

##### 3. Maintain

Once membrane potentials < −35 mV and/or action potentials with peak-to-peak amplitudes larger than 60 mV are obtained, the controller switches to ‘maintain’ mode, in which the position of electrode is kept constant with respect to neuron.

### Experimental procedures

#### Animals

*Aplysia Californica* weighing around 50 grams were obtained from the Rosenstiel School of Marine and Atmospheric Science, University of Miami, Florida. Animals were maintained in Instant Ocean™ artificial sea water (specific gravity of 1.25) at 13-16°C under 12:12 light-dark conditions.

#### Surgical Procedure

Animals were anesthetized by injecting 0.35 M Magnesium Chloride solution (7.7 g/L of MgCl_2_ and 3.6 g/L of HEPES) into the foot process for 5-10 min (equivalent to 30-35% of animal’s body weight). After the animal was distended and relaxed, it was pinned down on a dissecting dish and an incision was made along the entire length of the foot (from head to tail) to expose the internal organs and ganglia. The abdominal ganglion was identified and isolated with a fair amount of connective tissue. The ganglion was washed in artificial sea water (ASW, made with Instant Ocean™) three times and transferred to a petri-dish containing 2 ml of protease solution (1 mg/ml of 1 unit/mg Dispase II (Roche Diagnostics Corporation, Indianapolis, IN) in ASW). The ganglion was incubated in the protease solution at 34-35°C for 45 – 60 min to allow easy removal of connective tissue. The ganglion was then washed in ASW three times and placed in a Sylgard™ coated dish filled with ASW. The excessive connective tissue was used to pin down the ganglion using insect pins. Using surgical scissors (Item # 15000-08, Fine Science Tools, Foster City, CA) and fine forceps (Item # 500233, World Precision Instruments, Sarasota, FL), the connective tissue was carefully removed from the ganglion in order to expose the neurons for intracellular recording.

#### Intracellular recording and Analysis

Intracellular signals were recorded from abdominal ganglion neurons using an Electro 705 Intracellular amplifier (World Precision Instruments (WPI), Sarasota, FL) and the WPI 118 data acquisition system. Silver/silver-chloride pellet electrode (WPI, part # RC1T) was used as the reference electrode. The data acquisition system was accessed using the software, Labscribe™. The acquired raw signal was sampled at 10 KHz. All analyses were done in MATLAB™.

#### Force measurements and Analysis

Force generated by the electrothermal microactuator strips during one stroke has been estimated in our previous study [25]. Forces required to penetrate neurons in the abdominal ganglion of *Aplysia Californica* with a GP microelectrode at different penetration velocities were measured in order to evaluate the ability of electrothermal microactuators to impale cells. A GP microelectrode was wire-bonded to a custom-designed PCB, which was connected to a custom made connector board to interface with the intracellular amplifier. The PCB was attached to a connecting screw post and mounted on a precision 10 g load cell (Futek, LSB210, Irvine, CA). The load-cell with the microelectrode set-up was held on a hydraulic micromanipulator (FHC#50–12-1C, Bowdoin, ME) to move the electrode at different penetration speeds in randomized trials: 5 μm/s, 15 μm/s, 33.5 μm/s, 50 μm/s, 72 μm/s, 90 μm/s and 115 μm/s. For every trial, penetration of cell was confirmed from the voltage trace recorded simultaneously using the intracellular amplifier. Forces experienced by the electrode were acquired by the load cell at a sampling rate of 200 samples/sec. The acquired force data was smoothed using a 50 point moving average window and the peak forces were estimated using MATLAB™. Peak penetration forces were measured across the 7 speeds in n = 5 neurons and the variability in forces among neurons was evaluated using 1-factor ANOVA in OriginPro 8.5.1.

#### Measurements of electrical impedance of the electrode tip with increase in proximity to neuron (open-loop experiments)

To assess the change in electrical impedance of the tip as an electrode approaches a neuron, a single GP microelectrode was connected to a bond-pad on a custom PCB, which was attached to a custom-made connector board to interface with the intracellular amplifier. The PCB was mounted on a hydraulic micromanipulator (FHC#50–12-1C, Bowdoin, ME), which moved the microelectrode at 5 μm steps towards a neuron, until the electrode penetrated the cell (as confirmed by voltage recording). After each step, voltage responses to a set of five current pulses of 1 nA (50 msec ON, 2 sec OFF) were measured and the corresponding electrical impedance of the electrode tip were obtained in MATLAB™ using the algorithm in Fig. 3E. The final electrical impedance of the tip of the electrode at each position was calculated as the average of the five measurements.

#### Closed-loop control validation experiments

An Application Programming Interface (API) developed by iWorx Systems Inc. was used to enable MATLAB™ to communicate with the WPI 118 data acquisition system. A custom-built MATLAB™ routine continuously monitored the voltage recorded at the GP electrode tip. The routine also monitored the electrical impedance of the tip of the electrode at DC utilizing the same procedure applied for the open-loop experiments. The routine sent commands to WPI 118, which triggered the intracellular amplifier to inject current pulses through the electrode tip. Based on the two feedback variables, movement commands were sent from MATLAB™ to a custom-built Arduino-based waveform generator. The Arduino system generated the required activation waveforms for the hydraulic microdrive and subsequently the electrothermal MEMS microactuators to move the microelectrode towards a neuron, closing the feedback loop.

### In vivo recording

To assess the ability of a GP microelectrode to record intracellular activity in vivo, an anesthetized, adult Sprague-Dawley 500g male rat was used. All animal procedures were carried out with the approval of the Institute of Animal Care and Use Committee (IACUC) of Arizona State University, Tempe. The experiment was performed in accordance with the National Institute of Health (NIH) guide for the care and use of laboratory animals (1996). All efforts were made to minimize animal suffering.

#### Surgery, Implantation and data acquisition

The animal was induced using a mixture of (50 mg/ml) ketamine, (5 mg/ml) xylazine, and (1 mg/ml) acepromazine administered intramuscularly with a dosage of 0.1 ml/100 g body weight and maintained using 1-3% isoflurane. The anesthesia state of the animal was monitored closely throughout the experiment using the toe-pinch test. After mounting the animal on a stereotaxic frame (Kopf Instruments, Tujunga, CA, USA), a skin incision was performed to expose the skull. A 1 mm diameter hole with the center point at 3 mm posterior to bregma and 1.5 mm lateral to the midline was drilled using a burr drill. The dura and pia were carefully removed with a pair of micro scissors and the craniotomy was filled with phosphate buffered saline. Saline was applied to the exposed brain surface periodically to prevent it from becoming dry. The ground wire (silver) was wrapped around a metal screw implanted in the skull in the vicinity of the craniotomy. The GP microelectrode was implanted to a depth of 500 μm in the motor cortex using a conventional hydraulic micromanipulator. The electrical impedance of the tip of the electrode was measured before and after implantation to make sure that the tip was not clogged with debris. The electrode was moved in 2 μm steps at a speed of 3 μm/s using the hydraulic micromanipulator to seek neurons. Intracellular signals were acquired at a sampling rate of 20 KHz using the same set-up used for intracellular recordings from abdominal ganglion neurons of *Aplysia Californica*.

## Conclusions

In conclusion, we have developed and validated a miniaturized, robotic, MEMS based intracellular recording system comprising of glass-polysilicon penetrating microelectrodes, electrothermal microactuators and closed loop control for autonomous, high fidelity intracellular recordings from single neurons in the abdominal ganglion of *Aplysia Californica*. We have demonstrated the ability of glass-polysilicon microelectrodes to record intracellular signals that are comparable to recordings with conventional glass micropipettes. Further, we have shown that the success rate of penetration as well as the quality of recordings obtained with electrothermal MEMS based microactuators are similar to that of conventional stereotactic positioning systems using similar closed-loop control strategies. After careful in vivo validation in future experiments, this head-mountable system has the potential to record intracellular activity from a population of neurons simultaneously during behavior in awake animals due to its small form factor. This will impact several neurophysiological studies and enable discoveries on brain function and dysfunction. Further, this system will also enable the exciting possibility of repositioning electrodes in the event of loss of intracellular recordings.

## Acknowledgements

This research was supported by the NIH R21NS084492-01 grant. We would like to thank Vladislav Voziyanov for his help with the rodent craniotomy surgery, Dr. Arati Sridharan for her inputs during the fabrication of glass-polysilicon microelectrodes and Dr. Siddharth Kulasekaran for his help during the design of custom PC Boards. This paper describes objective technical results and analysis. Any subjective views or opinions that might be expressed in the paper do not necessarily represent the views of the U.S. Department of Energy or the United States Government. Sandia National Laboratories is a multi-mission laboratory managed and operated by National Technology and Engineering Solutions of Sandia, LLC, a wholly owned subsidiary of Honeywell International Inc., for the U.S. Department of Energy’s National Nuclear Security Administration under contract DE-NA0003525.

## Author contributions

SSK and JM were involved in the design, development, in vitro and in vivo testing of the glass polysilicon interfaces for intracellular recording and closed-loop controllers. SSK and JM were also involved in the integration and testing of all the microscale subsystems such as the intracellular penetrating microelectrodes, microscale actuators, control algorithms, packaging and interconnects. MSB, MO and JM were involved in the development of the microscale actuators and polysilicon microelectrodes using the SUMMiTV™ (Sandia’s Ultraplanar Multilayer MEMS Technology) process.

